# Modulation of C5a-C5aR1 signaling alters the dynamics of AD progression

**DOI:** 10.1101/2022.04.01.486759

**Authors:** Klebea Carvalho, Nicole D. Schartz, Gabriela Balderrama-Gutierrez, Heidi Y. Liang, Shu-Hui Chu, Purnika Selvan, Angela Gomez-Arboledas, Tiffany J. Petrisko, Maria I. Fonseca, Ali Mortazavi, Andrea J. Tenner

**Affiliations:** Department of Developmental & Cell Biology, University of California, Irvine, Irvine, CA 92697; Department of Molecular Biology & Biochemistry, University of California, Irvine, Irvine, CA 92697; Department of Neurobiology and Behavior, University of California Irvine, Irvine CA, USA; Department of Pathology and Laboratory Medicine, University of California, Irvine, School of Medicine, Irvine, CA, USA

**Author notes:** co-First author (Klebea Carvalho, Nicole D. Schartz). **Corresponding Author:** A.J. Tenner, University of California, Irvine, 3205 McGaugh Hall, Irvine, CA 92697-3900.

**Keywords:** complement, inflammation, Alzheimer’s disease, C5aR1, C5a, disease associated microglia, mouse model, therapeutic target

## Abstract

**Background:** The complement system is part of the innate immune system that clears pathogens and cellular debris. In the healthy brain, complement influences neurodevelopment and neurogenesis, synaptic pruning, clearance of neuronal blebs, recruitment of phagocytes, and protects from pathogens. However, excessive downstream complement activation that leads to generation of C5a, and C5a engagement with its receptor C5aR1, instigates a feed-forward loop of inflammation, injury, and neuronal death, making C5aR1 a potential therapeutic target for neuroinflammatory disorders. C5aR1 ablation in the Arctic (Arc) model of Alzheimer’s disease protects against cognitive decline and neuronal injury without altering amyloid plaque accumulation.

**Methods:** To elucidate the effects of C5a-C5aR1 signaling on AD pathology, we crossed Arc mice with a C5a overexpressing mouse (ArcC5a+) and tested hippocampal memory. RNA-seq was performed on hippocampus and cortex from Arc, ArcC5aR1KO, and ArcC5a+ mice at 2.7-10 months and age-matched controls to assess mechanisms involved in each system. Immunohistochemistry was used to probe for protein markers of microglia and astrocytes activation states.

**Results:** ArcC5a+ mice had accelerated cognitive decline compared to Arc. Deletion of C5ar1 delayed or prevented the expression of some, but not all, AD-associated genes in the hippocampus and a subset of pan-reactive and A1 reactive astrocyte genes, indicating a separation between genes induced by amyloid plaques alone and those influenced by C5a-C5aR1 signaling.. Biological processes associated with AD and AD mouse models, including inflammatory signaling, microglial cell activation, and astrocyte migration, were delayed in the ArcC5aR1KO hippocampus. Interestingly, C5a overexpression also delayed the increase of some AD-, complement-, and astrocyte-associated genes, suggesting the possible involvement of neuroprotective C5aR2. However, these pathways were enhanced in older ArcC5a+ mice compared to Arc. Immunohistochemistry confirmed that C5a-C5aR1 modulation in Arc mice delayed the increase in CD11c-positive microglia, while not affecting other pan-reactive microglial or astrocyte markers.

**Conclusion:** C5a-C5aR1 signaling in AD largely exerts its effects by enhancing microglial activation pathways that accelerate disease progression. While C5a may have neuroprotective effects via C5aR2, engagement of C5a with C5aR1 is detrimental in AD models. These data support specific pharmacological inhibition of C5aR1 as a potential therapeutic strategy to treat AD.

## Background

Alzheimer’s disease (AD) is the most common form of dementia and is characterized by the accumulation of extracellular amyloid beta (Aβ) plaques, hyperphosphorylated tau, synaptic loss, neuronal death, and ultimately cognitive decline (1). Evidence suggests that inflammatory pathways are induced by amyloid and tangle accumulation, contributing to AD pathologies, and exacerbating neuronal injury and cognitive decline (2-4).

The complement system is a powerful effector of the innate immune system that is activated via three distinct pathways, classical, lectin, and alternative, all of which converge on proteolytic cleavage of C3 into the chemoattractant C3a and opsonin C3b. C3b also forms part of the C5 convertase, which cleaves C5 into C5a, a potent pro-inflammatory chemoattractant, and C5b, the initiating molecule of the lytic membrane attack complex (MAC) (5). C1q, the recognition molecule of the classical pathway, in complex with the proteases C1r/C1s enables synaptic pruning as well as complement activation by fibrillar amyloid, which in the presence of C4, C2, C3, and C5 lead to the generation of C3a and C5a, both of which induce potent proinflammatory responses via their respective receptors C3aR and C5aR1 (6). In addition, administration of C5a to neurons *in vitro* is neurotoxic and enhances neurodegeneration (7, 8).

In two mouse models of AD, pharmacologic inhibition of the C5a receptor (C5aR1) with PMX205 reduced amyloid plaque accumulation, attenuated activation of microglia and astrocytes, and protected mice against hippocampal synaptic loss and cognitive decline (9). Inhibition or ablation of C5a-C5aR1 activity has also been demonstrated to increase survival and lower motor deficits in models of amyotrophic lateral sclerosis (ALS) (10, 11), reduce seizure susceptibility and inflammation in experimental epilepsy (12, 13), and accelerate functional recovery after spinal cord injury (14). These data are consistent with detrimental consequences of C5a-C5aR1 activity in neurodegenerative diseases, including AD, and that inhibition of C5a-C5aR1 signaling may slow or prevent the progression of these diseases. Importantly, C5aR1 antagonists have been shown to be nontoxic in human clinical trials and case studies (15-17). Genetic ablation of C5aR1 attenuated spatial memory decline and rescued neuronal integrity in the hippocampus in the Arctic mouse model of AD. This was accompanied by decreased inflammatory gene expression and enhanced expression of genes associated with debris clearance and phagocytosis in microglia, all supporting protective effects specific to C5aR1 inhibition (18). However, the molecular and cellular pathways involved remain to be discerned. Recent studies suggest that microglial activation can influence the reactive states of astrocytes (19). Whether C5aR1 ablation and the subsequent reduced microglia-mediated inflammation influences astrocytes in AD models remains to be determined.

C5aR1 expression is highly induced in AD mouse models (20). To further elucidate the role of C5a-C5aR1 signaling in amyloid-associated cognitive decline, and to expand on our previous behavioral studies examining the role of C5a-C5aR1 signaling on cognitive decline in the Arctic mouse, we crossed the Arctic mouse to a model that overexpresses C5a under the control of the GFAP promoter and tested spatial memory with the object location memory (OLM) test (18, 21). The dynamic transcriptomic and immunohistochemical changes due to C5a overexpression or C5aR1 ablation in Arctic mice were assessed using 6 mouse genotypes at 2.7, 5, 7, and 10 months of age. Since astrocytic and other cell-specific genes were missing from the analyses in our previous study, here, we microdissected the hippocampus and cortex of mice at different ages for bulk RNA-sequencing to identify region-, age-, and amyloid-specific transcriptomic changes. We identified more abundant changes in expression in hippocampi compared to cortices, correlating with higher amyloid pathology localization. We detected either decreased or delayed expression of AD-, disease-associated microglia (DAM), and reactive astrocyte-associated genes upon C5aR1 knockout. C5aR1 ablation also led to delayed expression of genes enriched for inflammation, microglial activation, phagocytosis, and cholesterol biosynthesis. Furthermore, we observed accelerated cognitive decline in the ArcC5a+, consistent with the hypothesis that C5a-C5aR1 signaling is deleterious in combination with amyloid pathology. Interestingly, C5a overexpression induced genes associated with synapse assembly, transcription, and GABAergic synapses, and may suggest parallel stimulation of C5aR1 and C5aR2, as a result of the high transgene driven expression of C5a, thus emphasizing the importance of specific targeted inhibition of C5aR1.

## Methods

### Animals

The Institutional Animal Care and Use Committee of University of California at Irvine approved all animal procedures, and experiments were performed according to the NIH Guide for the Care and Use of laboratory animals. Mice were grouped housed in ambient temperature and given access to food and water *ad libitum*. The Arctic48 mouse model of AD (hereafter referred to as Arc), which carries the human APP transgene with three mutations – the Indiana (V717F), the Swedish (K670 N + M671 L), and the Arctic (E22G), was generated on a C57BL6/J background and originally provided by Dr. Lennart Mucke (Gladstone Institute, San Francisco, CA). This hemizygous mouse model produces fibrillar plaques as early as 2 to 4 months of age (22). C5a-overexpressing mice, created by cloning the coding region for the C5a fragment of C5 plus a signal sequence into a construct containing the GFAP promoter to induce production and secretion of C5a as a function of GFAP induction (23), were crossed to Arc+/- mice generating wildtype (WT), Arc, C5a+ and ArcC5a+ mice. C5aR1KO mice, created by targeted deletion of the C5a receptor 1 gene (24) (originally provided by Dr. Rick Wetsel, Univ. of Texas Health Science Center, Houston), were crossed to Arc+/- mice to generate mice homozygous for C5aR1 deletion with and without the Arctic transgene. These mice were used to assess the effect of C5a overexpression on amyloid-associated cognitive decline, as well as the effects of C5a overexpression or C5aR1 ablation on microglial-, astrocytic-, and AD-associated gene expression and microglial and astrocytic pathology in WT and Arc mice from 2.7 months to 10 months of age. Both male and female mice were used in all experiments.

### Behavior

Object location memory (OLM) as previously described (18, 21). took place in dim lighting in testing arenas (37.3 × 30.8 × 21.6 cm) covered with approximately 1 cm of sawdust bedding. Briefly, mice were handled for 2 minutes each per day over 5 days in the testing room and were acclimated to the testing arena for 5 minutes per day over 6 days. The first day of habituation was used as an open field test to measure locomotion and anxiety-like behaviors. During the familiarization trial, two identical objects (e.g., 100 mL glass beakers, blue Duplo Legos, or opaque light bulbs) were placed in opposite and symmetrical locations in the testing arena and mice were allowed to freely explore the objects for 10 minutes. After a 24-hour delay, mice were tested for object location memory. One object was moved to the opposite end of the arena and exploration of both objects was tracked for two minutes. The object moved to a novel location was counterbalanced to control for side preference. To prevent olfactory distractions, objects were cleaned with 10% ethanol and bedding was stirred after each trial (different bedding was used for males and females). Trials were recorded by mounted cameras from above and object exploration was scored manually by 2 blinded experimenters using stopwatches. Discrimination indices were calculated with the formula: [(time spent with moved object - time spent with unmoved object)/time spent with both objects) x 100] to obtain a percent (%) discrimination index (DI) (21). Mice were removed from the analysis if they spent less than 1 second/minute with the objects during training and/or testing or if the performance of mice was ± 2 standard deviations from the mean.

### Immunohistochemistry (IHC)

Mice were deeply anesthetized with isoflurane and perfused transcardially with cold phosphate buffered saline (PBS; 137 mM NaCl, 2.7 mM KCl, 4.3 mM Na_2_HPO_4_, 1.47 mM KH_2_PO_4_, pH 7.4). Half brains were quickly dissected and fixed in 4% paraformaldehyde for 24 hours, then stored in PBS with 0.02% sodium azide at 4°C. 30-40 µm coronal sections were obtained.

Sections were incubated in blocking solution (2% BSA plus 5% normal goat serum or 10% normal goat serum, and 0.1% Tx PBS) for 1 hour on a shaker at RT. Tissues were then incubated with primary antibody diluted in blocking solution overnight on a shaker at 4°C. Primary antibodies used were rabbit anti-Iba1 (1:1000, Wako #019-19741), rat anti-CD68 (1:700 Biolegend #137001), rat anti-CD11b (Biorad #MCA74G), hamster anti-CD11c (1:400, Biorad #MCA1369), rat anti-Lamp1 (1:500, Abcam #ab25245), rat anti-mouse C3 (1:50, Hycult #HM1045), rabbit anti-GFAP (1:2900, Dako #Z0334). Alexa Fluor secondary antibodies were diluted 1:500 in blocking solution and included 568 goat anti-Armenian hamster (Abcam, # ab175716), 555 goat anti-rat (Invitrogen #A21434), 488 goat anti-rat (Invitrogen #A21212), 488 goat anti-rabbit (Invitrogen #A-11070), and 647 goat anti-rabbit (Invitrogen #A21244). To counterstain fibrillar plaques with Thioflavin-S (Thio-S) sections were incubated in 0.1% or 0.5% ThioS for 10 min after secondary antibody. To counterstain fibrillar plaques with Amylo-Glo (1:100 in PBS, Biosensis #TR-300-AG), tissues were incubated for 10 min before the first blocking step, washed in PBS for 5 min, and quickly rinsed with MilliQ H_2_O before continuing with the IHC protocol. Sections were mounted and coverslipped with Vectashield (VECTOR). Low magnification images (10X) were acquired using ZEISS Axio Scan.Z1 Digital Slide Scanner. The mean areas of C3, GFAP, CD11b, CD11c, CD68, Iba1, and plaques were quantified in the hippocampal regions CA1, CA3, and DG, as well as the entire hippocampus and cortex using the Surfaces feature of Imaris x64 (version 9.5.0). Confocal images were acquired at 20 x using a Leica SP8 confocal microscope. Z stacks of 1um step interval within a depth of 30 um were obtained per area of interest, and volume was analyzed with Imaris. Quantitative comparisons between groups were carried out on comparable sections of each animal processed at the same time with same batches of solutions, unless otherwise noted.

### Enzyme-linked immunosorbent assay

To confirm that mice expressing the C5aGFAP+/- transgene produce higher quantities of C5a protein, we performed enzyme-linked immunosorbent assay (ELISA) on pulverized hippocampus and cortex and on plasma taken from WT, Arc, C5a+, and ArcC5a+ mice at 5, 7, and 10 months. Pulverized cortex and hippocampal samples were homogenized on ice using a pestle pellet motor in 10X volume of homogenization buffer containing PhosphoSTOP (Roche) and cOmplete Mini Protease Inhibitor Cocktail (Roche) in TPER. Samples were then centrifuged at 14,000 rpm for 30 min at 4°C and supernatant was collected and stored at -80°C. C5a concentration was determined with the R&D Systems Mouse Complement Component C5a DuoSet ELISA (DY2150) according to the manufacturer’s instructions, in triplicate. Optical density was read on SpectraMax Plus 384 using and SoftMax Pro 7.1 (Molecular Devices, San Jose, CA). Standard curve was fitted to 4-parameter logistic expression. The total protein concentration was assessed using the Pierce BCA Protein Assay Kit (Thermo Scientific). Any plates lacking a full standard curve were normalized to the previous plate while C5a levels were averaged to generate one value per animal for samples repeated in more than one ELISA experiment. To assess whether the C5aGFAP transgene resulted in peripheral effects, plasma levels of C5a were measured. Blood collected via cardiac puncture was mixed immediately with 10mM EDTA on ice. After centrifugation (5,000 rpm for 10 min, 4°C), plasma was collected and diluted 1:80 in reagent diluent for ELISA. When plasma from FVB/NJ C5-deficient mice (25) were assayed no C5a was detected, validating our C5a ELISA.

### RNA extraction

Mice were perfused with PBS and dissected cortex and hippocampus were stored at - 80°C prior to RNA extraction. Hippocampus and cortex were lysed separately in RLT (Qiagen # 80204) buffer with 1% β-Mercaptoethanol by the QIAGEN TissueLyser. Total RNA of each tissue was extracted using the QIAGEN RNeasy mini kit and QIAcube (Qiagen # 80204 and # 9002864) and quantified using the NanoDrop ND-1000 spectrophotometer. The RNA integrity number (RIN) was assessed using the Agilent 2100 Bioanalyzer and the samples with RIN > 8.0 were used for library preparation.

### RNA-seq library preparation

Bulk sequencing experiments were conducted utilizing 5-10 mice per genotype per age (with approximately equal numbers of males and females). RNA-seq libraries were built following the Smart-seq2 protocol using the Nextera library preparation kit (26). In brief, polyadenylated RNA is reverse transcribed and a template-switching oligo (TSO) is added, which carries 2 riboguanosines and a modified guanine to induce a locked nucleic acid (LNA). cDNA was then amplified, and the resulting fragments were tagmented. Fragments between 150 and 600 nucleotides were finally selected using Ampure XP beads. The quality of all libraries was assessed using the Agilent 2100 Bioanalyzer. Bulk libraries were sequenced using the NextSeq 500 (Illumina) obtaining at least 10 million reads per RNA-seq sample. Resulting Fastq files and data matrices were deposited in GEO with the accession ID: GSE197591.

### RNA-seq processing and data analysis

Paired-end RNA-seq reads were aligned to mm10 reference genome and annotated with Gencode v21 transcriptome using STAR v.5.1 (27). Gene expression was calculated using RSEM v1.2.25 (28). Possible noise introduced by batch effects was corrected by the R (version 3.6.2) package Combat-seq (29). Data was then normalized using edgeR trimmed mean of M-values (TMM) function (30). TPM was calculated utilizing a custom script and ComBat-seq batch corrected counts as input. Two statistical outliers were removed based on dimensionality reduction by PCA and Pearson correlation coefficient (31). A total of 372 RNA-seq data sets were generated. We also provide an explorable data viewer that presents all generated RNA-seq data (http://crick.bio.uci.edu:3838/ArcticMonday/).

### Differential expression analysis

The R package maSigPro (version 1.58.0) was used to identify gene expression changes over time allowing for k-means clustering of genes that present similar patterns of expression during the time course (32). In addition, edgeR (version 3.28.1) was used to identify genes differentially expressed between selected ages and genotypes (30) using a false discovery rate (FDR) of 1% and an alpha of 0.05. We utilized expression as TMM normalized counts represented as a count per million (CPM) matrix in both aforementioned packages.

### Gene ontology and pathway analysis

Gene ontology enrichment analysis was performed using Metascape and DAVID online tools computing gene-set overlaps between pathways and biological processes, that were selected based on p-values smaller than 0.05 (33, 34).

### Statistical Analyses

Unless otherwise stated, all statistical analyses were performed with GraphPad Prism (V9.3.1). When appropriate comparisons were performed with two-way ANOVA and Tukey’s post hoc test.

## Results

### C5a-C5aR1 signaling accelerates hippocampal-dependent memory deficits in Arctic mice

We previously reported that ablation of C5aR1 in Arc mice prevents memory deficits assessed with the OLM test at 10 months of age (18), suggesting that binding of C5a to C5aR1 induces responses that contribute to the cognitive decline observed in this model. Since elevated C5a production could result in greater engagement of C5a with C5aR1, we assessed whether transgene-driven generation of the C5a fragment accelerates the effects of this receptor signaling. We first confirmed that the C5a+ transgene resulted in elevated levels of C5a in the cortex and hippocampus in both WT and Arctic mice without increasing plasma levels (**Figure S1**), as expected since it is under the control of the GFAP promoter. Arc and ArcC5a+ mice and their WT littermate controls were tested at 7 months to determine if overexpression of C5a in brain would accelerate memory decline in the presence or absence of amyloid pathology. Analysis of open field revealed that there were no abnormalities in locomotion or anxiety-like behavior (**Figure S2A-C**). All mice had similar total exploration times of both objects (**Figure S2D**) and none showed a preference for either object position during OLM training (**Figure S2E)**. At 7 months of age, Arc mice did not show a deficit in object location memory compared to WT (**Figure 1**). However, while no deficit was detected in the C5a overexpressing wildtype, the DI of ArcC5a+ mice was 6.2%, indicating significantly diminished cognitive performance compared to Arc mice (41.9%) (**Figure 1B**). These data suggest that higher levels of C5a act in collaboration with co-existing amyloid pathology (which also results in induced expression of C5aR1) to accelerate cognitive decline. Taken together, and with the data published in 2017 (18), these findings are consistent with the hypothesis that chronic stimulation of the C5a-C5aR1 accelerates memory decline in Arc mice, whereas ablation of C5aR1 protects against it.

**Figure 1:**
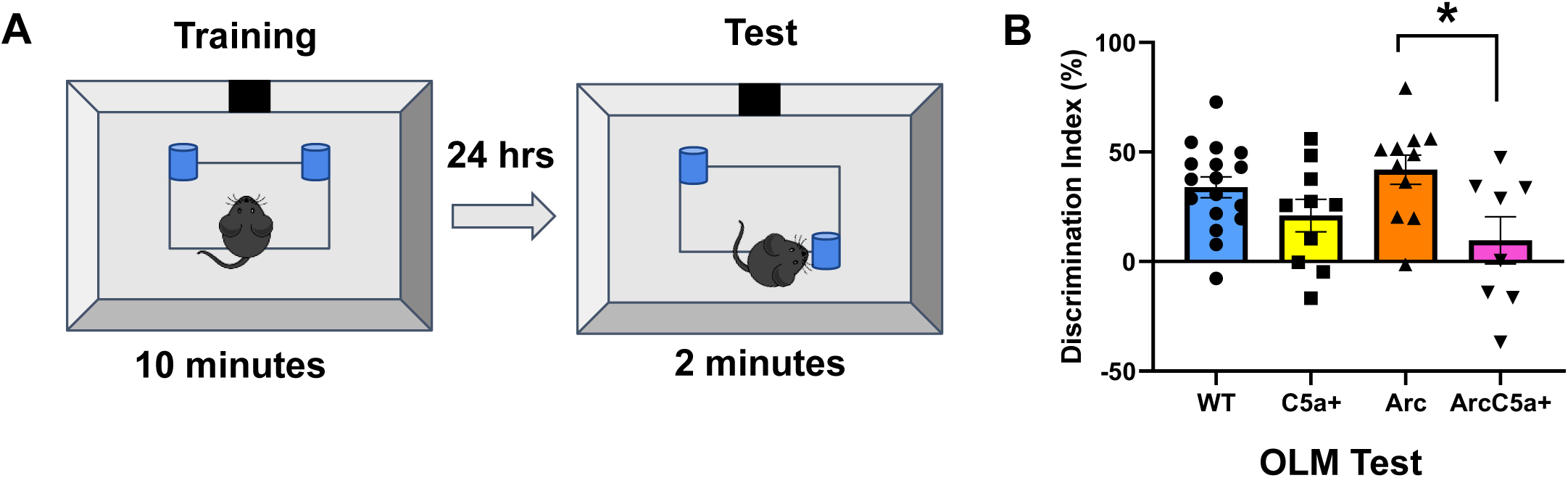
Overproduction of C5a accelerates memory decline in Arctic mice at 7 months. **(A)** Overview of experimental design for object location memory (OLM) test. (**B**) Discrimination index (%) was calculated for the OLM test. Data shown as Mean ± SEM. * *p* < 0.05. Two-way ANOVA with Tukey’s post hoc test. N = 17 (WT), 10 (C5a+), 11 (Arc), 8 (ArcC5a+).

Neuronal loss and cognitive decline are common features of Alzheimer’s disease (1). Therefore, to determine if hippocampal-dependent cognitive decline in Arc and ArcC5a+ mice paralleled neuronal injury in the hippocampus, we assessed levels of dystrophic neurites (Lamp1) (**Figure S3A-B, S3E**). While ArcC5aR1KO mice had lower levels of Lamp1-positive dystrophic neurites compared to Arc in the CA1, CA3, and DG of the hippocampus and in the cortex at 5 mo (and in the DG at 7 mo) (**Figure S3E**) by 10 mo the levels of LAMP1 were comparable in Arc and ArcC5aR1KO. ArcC5a+ had similar levels of Lamp1 as Arc, with a reduction only in the DG at 7 months (**Figure S3E**). To determine if the toxic response to fibrillar plaques was altered with the deletion of C5aR1 or overexpression of C5a, we compared the relative levels of Lamp1 to Amyloglo (AG) levels in Arc, ArcC5aRKO, and ArcC5a+ (**Figures S3C-D & S3F-G**). While there were minimal reductions of AG at 7 months in the ArcC5aR1KO CA3 and DG hippocampal regions (**Figure S3F**), the ratio of Lamp1 to AG was significantly reduced in ArcC5aR1KO and ArcC5a+ at 10 months (**Figure S2G**). These data suggest that while there may not be lasting changes in fibrillar plaque accumulation in ArcC5aR1KO or ArcC5a+ mice, the toxic response to these plaques is reduced.

### Distinct subsets of genes are affected by C5ar1 knockout or C5a overexpression

To explore molecular basis of C5a-C5aR1 signaling, we dissected 372 cortices and hippocampi of mice representing six distinct genotypes (WT, C5aR1KO, C5a+, Arc, ArcC5aR1KO, and ArcC5a+) throughout disease progression at 2.7, 5, 7, and 10 months of age in order to build RNA-seq libraries and thus identify transcriptome changes in different cohorts (**Figure 2A**). UMAP dimensionality reduction showed clear tissue-specific clustering characterized by the separation of cortices from hippocampi samples on UMAP 1. UMAP 2 showed a subtle separation of tissues from younger mice located at the top and tissues of older mice located towards the bottom of the axis in hippocampus samples, and left to right in cortex samples, highlighted by the gradient of colors with age from light to dark, respectively (**Figure 2B**). To determine if the Arctic model has sex-specific effects, we used edgeR (30) to identify differentially expressed genes between male and female Arc mice. Genes up-regulated in females were mainly those known to be sexually dysmorphic genes (35), such as Xist and Tsix, which are located on the X-chromosome, whereas males presented upregulation of genes known to be male-specific, such as Eif2s3y and Ddx3y, which are located on the Y-Chromosome (**Figures S4A-B**). Therefore, differential expression analyses showed lack of sex-specific changes in the hippocampus and cortex of Arc mice at all ages.

**Figure 2:**
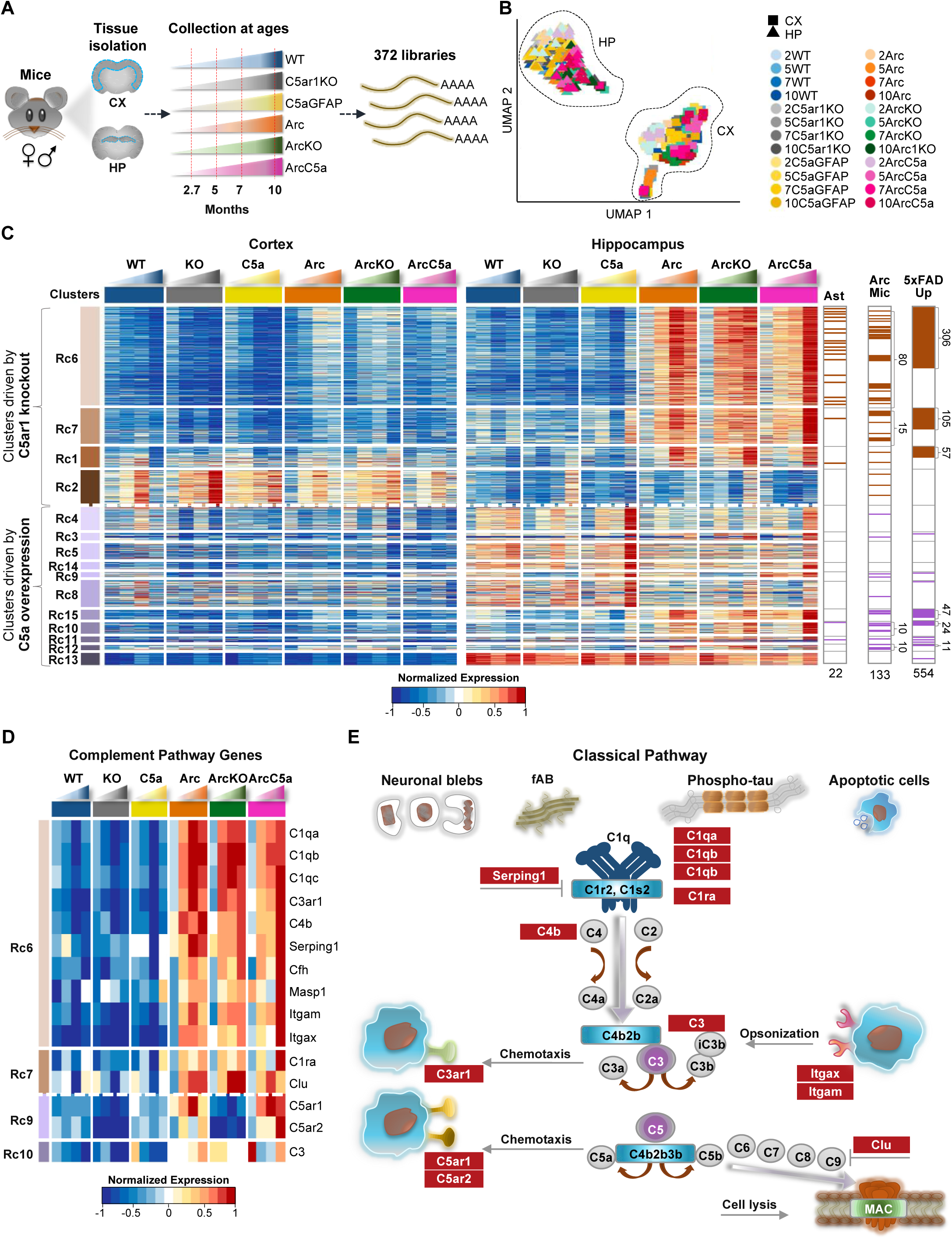
Distinct subsets of genes are affected by C5ar1 knockout or C5a overexpression. (**A**) Schematic diagram of experimental design highlighting 372 RNA-seq samples processed from cortices and hippocampi of 6 mice genotypes during disease progression at 2.7, 5, 7, and 10 months of age. (**B**) UMAP embedding representation of 372 RNA-seq samples represented in panel A. Samples from WT mice are represented in shades of blue, from the C5ar1KO cohort in shades of gray, from the C5aGFAP cohort in shades of yellow, from the Arctic cohort in shades of orange, from the ArcC5ar1KO cohort in shades of green, and samples from the ArcC5aGFAP cohort in shades of pink. (**C**) Heatmap of 1,763 genes with dynamic temporal profiles identified by maSigPro clustering (alpha < 0.05, FDR < 0.05%). Each column represents the average expression for a time point and each row represents a gene. Each cluster represents a subset of genes that show a similar pattern of expression along the time course. Clusters shown in brown gradient and lilac gradient represent genes whose expression changes are driven by C5ar1KO or C5a overexpression, respectively. RNA-seq data (TPM) is row-mean normalized. Astrocyte-associated genes (Ast), Arctic mice isolated microglia-associated genes (Arc Mic), and genes upregulated with pathology progression in the 5xFAD mice (5xFAD Up) present in each cluster are shown. (**D**) Heatmap of 15 genes of the complement pathway that were present in maSigPro identified clusters. RNA-seq data (TPM) is row-mean normalized. (**E**) Representative diagram of the Classical pathway activation of the complement cascade, highlighting complement-associated genes present in the maSigPro identified clusters (as seen in panel **D**). N = 4-10 mice/genotype/age

Using maSigPro (**Figure S5A**) (32), we identified 1,763 genes whose expressions varied in a time-specific fashion. These genes grouped into 15 RNA clusters with distinct patterns of expression (**Figure 2C**). RNA cluster (Rc)6, Rc7, Rc1, and Rc2 represent genes whose expression changes were mainly driven by the knockout of C5ar1 and are represented in shades of brown. Rc4, Rc3, Rc5, and Rc8 to Rc15 represent genes whose expression changed mainly in response to C5a overexpression and thus are represented in shades of lilac. Each cluster contains distinct subsets of complement pathway (CP) genes, as well as distinct subsets of astrocyte-associated (Ast) (19), Arc microglia-associated (Arc Mic) (18), and AD-associated genes elevated in the 5xFAD mouse model with pathology (5xFAD up) (36) (**Figure 2C and S5B-C**). Expression of the C5ar1 gene was induced in Arc relative to WT as early as 2.7 mo and increased more dramatically at 5 and 7 mo, while was lowest (essentially non detectable) in the C5aR1KO and ArcC5aR1KO cohorts, confirming its genetic ablation (**Figure 2D**). Expression of C5ar2 was also decreased in the C5aR1KO and ArcC5aR1KO cohorts, consistent with previous studies showing a correspondence in expression of those genes (37, 38), but strongly induced in the ArcC5a+ particularly at 10 mo. The ArcC5a+- and ArcC5aR1KO-dependent expression changes in Rc6, Rc7, and Rc1 were more abundant in the hippocampus compared to cortex, correlating with the relative density of amyloid pathology in the Arc mice (**Figure S6C**), and thus suggesting synergy/interaction between the presence of amyloid plaques and C5aR1 signaling.

Expression of genes of some of the early complement pathway components were induced in Arc mice with a peak at 7 months of age and suppressed in the absence of C5ar1. Notably, cluster Rc6 contained most of the complement genes identified using maSigPro including C1qa, C1qb, C1qc (known to be coordinately expressed) (39, 40), and C4b (designation for the mouse C4 gene), as well as the receptor for the C3a activation fragment, C3ar1, and the inhibitor Serping1 (inhibitor of C1, as well as bradykinin formation), all of which showed higher expression in the Arc mice at 5 months and increasing with age (**Figure 2D**). The earliest upregulated complement related genes in Arc relative to WT were C5aR1, C4b, Serping1 and C3 as well as C1q, and C3ar1 at 2.7 mo of age. While C1q was robustly upregulated in all Arc genotypes, consistent with the rapid increase in multiple other injury models, increases in expression of the genes for C4 (required for synaptic pruning), Factor H and Serping1 were slower in the ArcC5aR1KO as seen in the previous analysis of isolated microglia in this model (18). Interestingly, delayed increases in these components were also seen in ArcC5a+, but by 10 months expression levels of these genes reached or surpassed that seen in the Arctic.

Additional changes in expression of complement genes that were driven by transgene overproduction of C5a. Complement genes Masp1 and C1ra, regulators Cfh and Serping1, and the α chains Itgam (CD11b) and Itgax (CD11c) of CR3 and CR4, respectively, were upregulated early by the presence of the Arctic APP transgene but showed increased expression in the ArcC5a+ hippocampus at 10 months (**Figure 2D**). Thus, the expression of individual complement genes may be orchestrated to fit the local environment with the knockout of C5aR1 and the overexpression of C5a affecting the level and time of induction of genes that act in multiple arms of the complement cascade, including complement factors, receptors, and inhibitors (**Figure 2E)**.

### C5aR1 ablation may delay microglial switch from disease mitigating to disease enhancing

To understand the effects of C5a-C5ar1 signaling on gene expression in AD, changes in expression of genes found in the current bulk RNA analysis were compared to our previous reports of microglia isolated from combined cortex and hippocampus from Arc and ArcC5aR1KO at the same ages (18), to those known to be associated with AD pathology progression in the 5xFAD mice model (36), and to human AD genes (41-44) (**Figures 2C, right & S5B**). Most of the genes upregulated in previously assessed microglia isolated from the Arc mice were part of clusters Rc6 (80 genes) and Rc7 (15 genes), which are linked to inflammation and cholesterol metabolism, respectively (**Figure 2C**). Similarly, most of the genes upregulated in the brain of the 5xFAD mouse model were part of Rc6 (306 genes) and Rc7 (105 genes). While many genes were upregulated with accumulation of amyloid regardless of C5a-C5aR1 status, AD genes Cd33, Trem2, Tyrobp and Cst7 **(Figure 3A)** showed increased expression in the Arc cohort with a peak at 7 months of age. However, the ArcC5aR1KO the expression increased at a lower rate and did not peak until 10 months of age. Inpp5d, S100a6, and Stat3 showed lower expression at all ages in the ArcC5aR1KO (**Figure 3A and S5B)**. It has been shown that Inpp5d expression increases in early stages of AD in Japanese patients, is positively correlated with plaque load in LOAD (45) and increases with progression of disease in the 5xFAD mouse model (46). Similarly, S100a6 is upregulated in patients with AD, as well as AD mouse models (47). Stat3 has been associated with worsening cognitive decline in the 5xFAD mice (48). Thus, C5ar1 deletion either delayed or prevented the expression of several genes upregulated in the Arctic and 5xFAD mouse models and human AD. In summary, eliminating C5ar1 in the Arctic AD model results in a decrease or delay of expression of many, but not all, reactive glial genes associated with AD in human or mouse models, indicating a separation between those genes induced by amyloid plaques (or other sources of “damage”) and those requiring the C5a-C5aR1 signaling.

**Figure 3:**
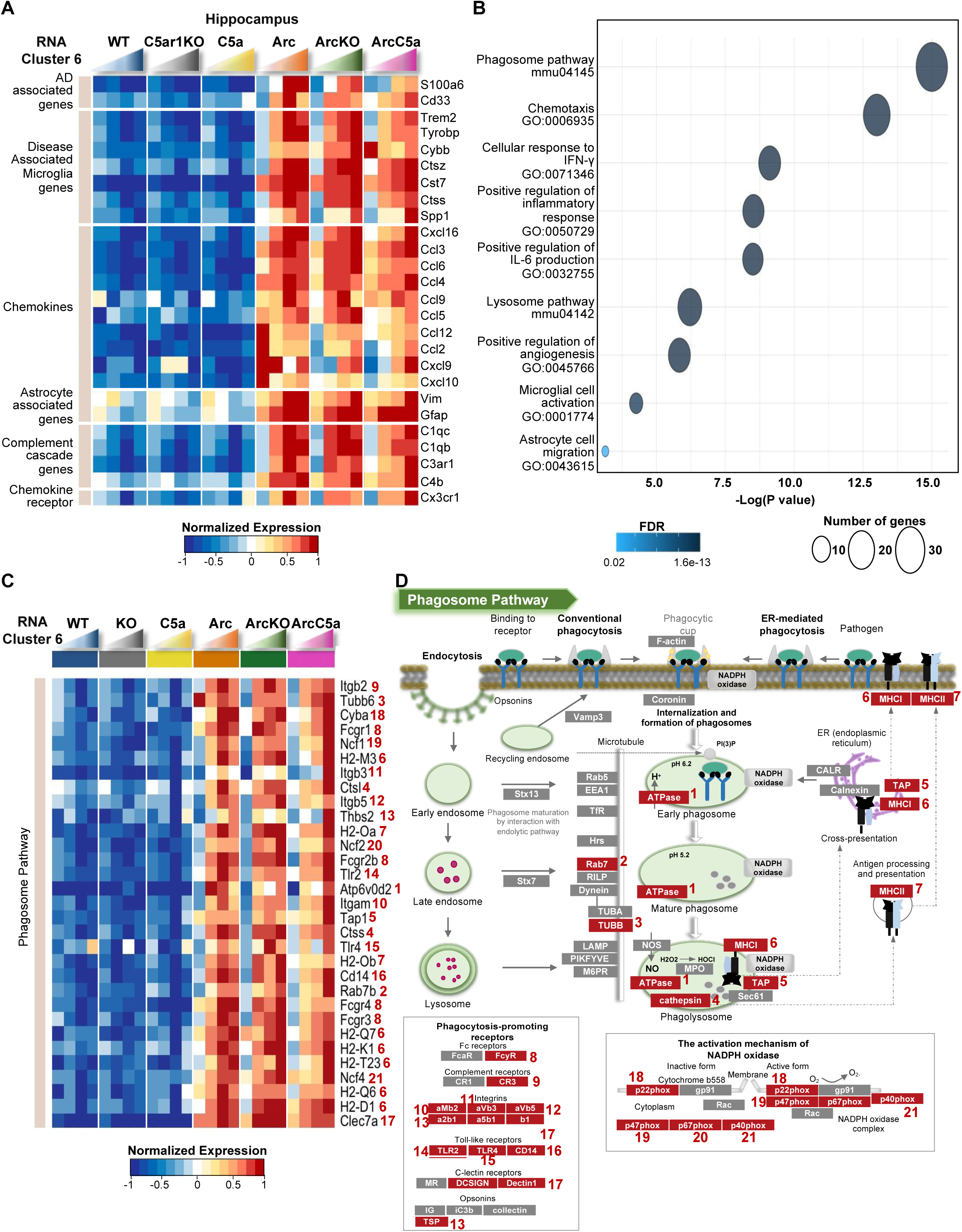
Delay and/or reduction of inflammation- and DAM-associated genes in the ArcticC5aR1KO mice. (**A**) Heatmap of selected genes present in RNA cluster 6. Genes were clustered based on AD study findings and molecular functions. RNA-seq data (TPM) is row-mean normalized. (**B**) Gene ontology (GO) and pathway enrichment analyses of genes present in Rc6. (**C**) Heatmap of genes enriched in Phagosome pathway (mmu04145). Numbers in front of gene names correspond to protein they encode, as seen in panel **D**. (**D**) Representative diagram of the Phagosome pathway (mmu04145) with select protein encoded by genes present in the panel **C** heatmap highlighted in red. N = 4-10 mice/genotype/age

### C5a-C5aR1 signaling influences astrocyte activation

While many of the above genes are considered predominantly microglial genes, given the evidence for altered astrocyte states in AD (19, 49, 50), we investigated the changes in expression of astrocyte-associated genes. Rc6 contained 18 astrocyte-associated genes, including pan-reactive astrocyte genes Lcn2, Osmr, Cd44, Vim, and Serpina3n and A1 reactive astrocyte genes Ggta1, H2-T23, Srgn, and Serping1 (19), whose expression was higher in the Arc cohort. Ablation of C5ar1 reduced or delayed the activation of those genes (**Figure S5C**), suggesting that C5ar1 deletion may reduce the activation of neurotoxic reactive astrocytes. Interestingly, overexpression of C5a led to increased expression of these pan-reactive and A1 genes at 10 months suggesting that an increase in C5a may lead to activation of primed reactive astrocytes, either directly or indirectly. GFAP (pan-reactive) and Aspg and Psmb8 (A1) were highly upregulated in the presence of amyloid plaques regardless of C5a-C5aR1 manipulation, indicating a subset of astrocyte genes that are responsive to insult, but not influenced by C5aR1 signaling (**Figure S5C)**. Assessing the microglial mediators of astrocyte activation, TNFα, IL1α and C1q (19, 51), we found that while IL1α was elevated at 5 months in the ArcC5aR1KO relative to Arc and ArcC5a+, TNFα expression was substantially delayed in ArcC5ar1KO, and C1qa-c were transcribed with the same temporal increase in response to plaque accumulation for all Arc genotypes (**Figures 2D, S5B, S5D**). These findings indicate a complexity of astrocyte response mediators that are induced by different insults or injury and demonstrate that expression levels of some powerful mediators can be modulated by C5a-C5aR1 signaling.

### Gene ontology analysis of Rc6 revealed reduced expression of inflammation- and DAM-associated genes in the ArcC5aR1KO mice

We performed gene ontology analysis to identify relevant biological processes and pathways associated with differentially expressed genes present in each maSigPro cluster. Expression of genes in Rc6 was higher in Arc mice and peaked in the hippocampus of the Arc cohort at 7 months. However, in the ArcC5a+ cohort, expression did not increase until 10 months of age. Rc6 was enriched for DAM genes, such as Trem2, Tyrobp, Cybb (gp91 of the NADPH oxidase), Cst7, Ctss, and Spp1 (**Figures 3A & 3C**). Rc6 also contained inflammatory chemokine genes such as Ccl12, Ccl2, and Cxcl10 that were highly expressed in the Arc at 2.7 mo before dropping and then slowly increasing. Neither ArcC5aR1KO nor ArcC5a+ expressed these genes at 2.7 mo prior to plaques deposition. Additionally, gene ontology and pathway analysis showed enrichment for biological processes that are significant in the AD context, for instance regulation of inflammatory signaling including IFN-γ and IL-6 production, as well as microglial cell activation and lysosome pathway, all of which were delayed in the ArcticC5aR1KO cohort (**Figure 3B**). Our results suggested that the ablation of C5ar1 delayed disease enhancing activation genes without diminishing the disease mitigating response to injury (**Figure 3C**).

### Gene ontology analysis of Rc7 revealed reduced expression of genes associated with lipid and cholesterol biosynthesis and metabolism in the ArcC5aR1KO mice and increased expression of genes of those genes in the ArcC5a+ mice

Rc7 mainly consisted of genes whose expression was increased in the Arc peaking at 7 months. While expression in the ArcC5aR1KO remained higher than C5aR1KO controls, it did not increase uniformly with age. Interestingly, genes in Rc7 were highly upregulated in the ArcC5a+ mice at 10 months compared to earlier ages and Arc and ArcC5aR1KO (**Figure 4A**). Rc7 also contained genes associated with inflammatory response, such as Csf1, Icam1, Tnfsf1b, Akna, Sbno2, and Nrros and cytokine signaling, such as Jak3 and receptor Tgfbr2, whose expressions were reduced in the absence of C5ar1. Gene ontology and pathway analysis showed enrichment for lipid metabolic processes, inflammatory response, and cholesterol biosynthesis, suggesting that those processes were induced by amyloid plaque challenge but reduced upon C5ar1 deletion in the Arctic model at 7 months while upregulated at 10 months when C5a is overexpressed (**Figures 4A-B**). Cholesterol cannot cross the blood brain barrier, and thus almost all brain cholesterol is synthesized locally. Interestingly, many of these genes are also upregulated in the C5a+ in the absence of the APP transgene at 10 mo of age. Our results suggest that ablation of C5ar1 reduces brain cholesterol metabolic processes or the need for such processes, and that cholesterol metabolism in the brain may be partially influenced by high concentrations of C5a (**Figure 4C**).

**Figure 4:**
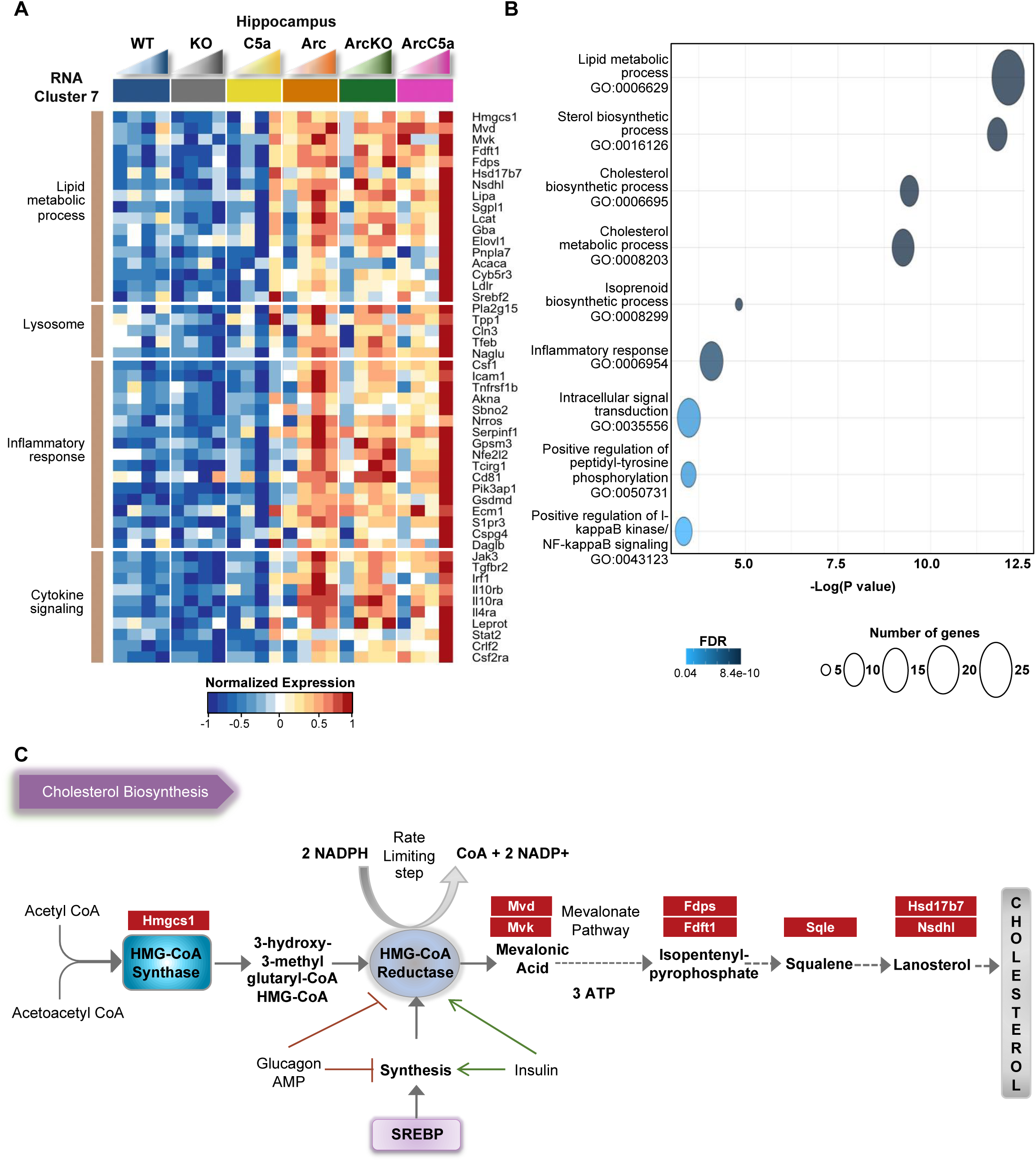
Reduced cholesterol biosynthesis-associated genes in the ArcticC5aR1KO mice. (**A**) Heatmap of selected genes present in RNA cluster 7. Genes were clustered based on biological processes and molecular functions. RNA-seq data (TPM) is row-mean normalized. **(B)** Gene ontology (GO) and pathway enrichment analyses of genes present in Rc7. (**C**) Representative diagram of the Cholesterol biosynthesis pathway with genes present in the panel A heatmap highlighted in red. N = 4-10 mice/genotype/age

### C5a overexpression leads to increased expression of genes associated with synapse transmission and assembly

C5a has been shown to promote neuronal damage in vitro (8), although it plays a positive role in neurogenesis during early development (52). We thus sought to explore effects of C5a overexpression in our mice in the presence and absence of amyloid pathology, given the accelerated behavior deficiency in the ArcC5a+ mice relative to the Arc at 10 months. In clusters 3 and 4, there was a general decrease with age in the Arc mice relative to the wildtype controls, except for a marked increase in ArcC5a+ mice at 10 mo but an even greater increase in gene expression in the C5a+ mice at 10 months in the absence of the amyloid transgene (**Figure 5A**). C5a overexpression promoted pathways associated with GABAergic synapses, ion transport, synapse assembly, axogenesis, and transcription (**Figure 5B**), more so in the absence of the Arc transgene. Specifically, C5a overexpression was associated with an increase of genes that regulate GABAergic synapse processes (**Figure 5C**). It has been shown that alterations of GABAergic neurotransmission may contribute to AD pathology (53, 54). Therefore, C5a overexpression might contribute to alterations of GABAergic transmissions independently of amyloid pathology. Whether these aberrant synaptic processes are associated with the acceleration of cognitive decline in ArcC5a+ mice is yet to be explored.

**Figure 5:**
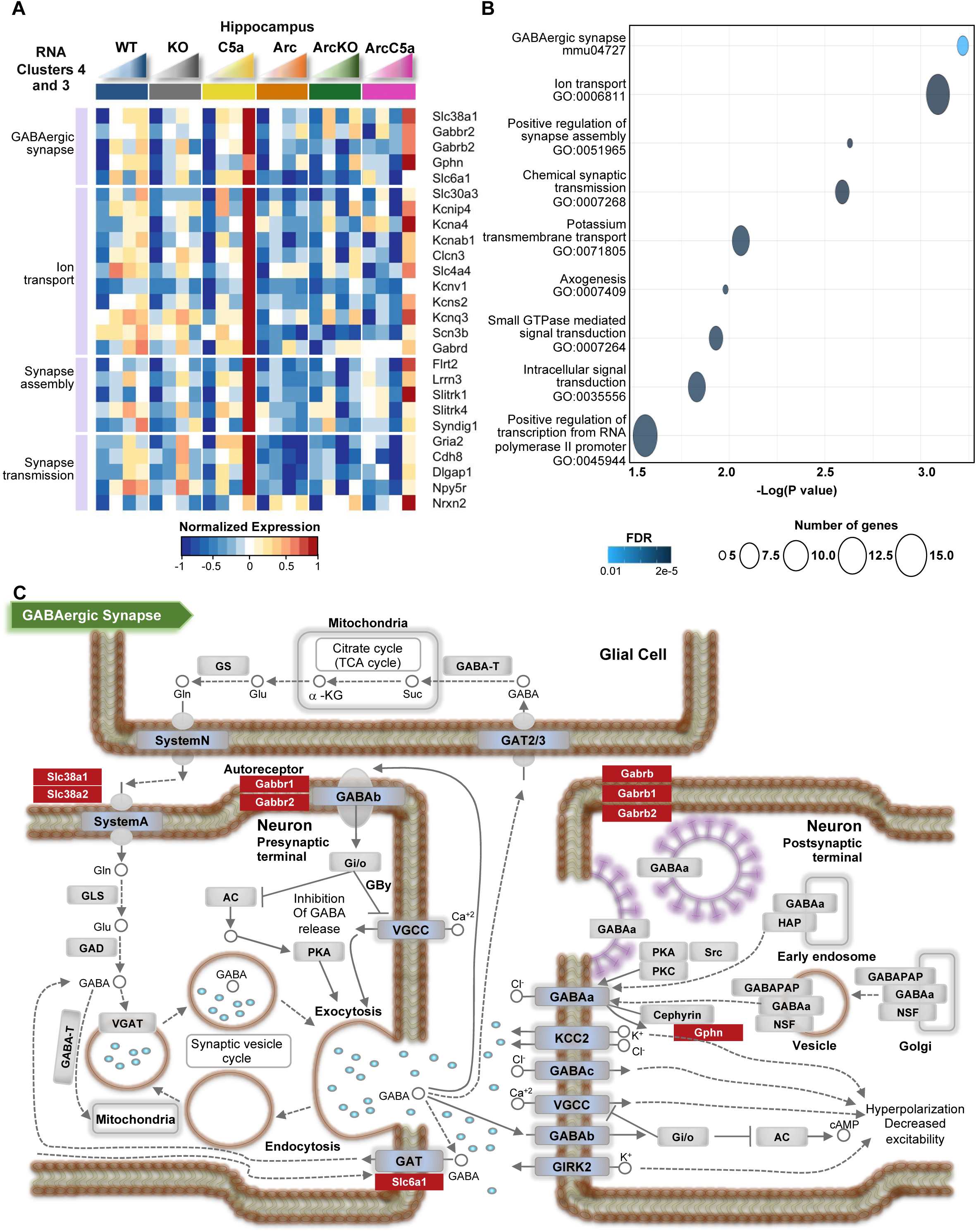
C5a overexpression leads to increase in expression of genes associated with synapse transmission and assembly. (**A**) Heatmap of selected genes present in RNA clusters 3 and 4. Genes were clustered based on biological processes. RNA-seq data (TPM) is row-mean normalized. (**B**) Gene ontology (GO) and pathway enrichment analyses of genes present in Rc6. (**C**) Representative diagram of the GABAergic synapse pathway with select genes present in the panel A heatmap highlighted in red. N = 4-10 mice/genotype/age

### Modulation of microglia activation marker CD11c in Arctic mice by C5a signaling

Amyloid pathology in humans and animal models can activate microglia to an inflammatory and disease-enhancing state (55). We observed evidence of a delay in some phagosome and inflammatory pathways driven by elimination of C5aR1 and an exacerbation driven by overexpression of C5a at 10 mo in the RNAseq data. To determine the effects of C5aR1 ablation or C5a overexpression on amyloid-associated microglial protein expression, we stained ArcC5aR1KO and ArcC5a+ tissue with antibodies for CD11b and CD11c and compared levels to Arctic at 5, 7, and 10 months (**Figure 6**). The hippocampus and cortex were assessed separately as amyloid deposition and glial activation are seen first, and to a larger extent, in the hippocampus compared to the cortex in the Arc model (**Figures S6A-B**). As previously noted, amyloid deposition in the hippocampus remained essentially unchanged with deletion of C5aR1 or overexpression of C5a, with only small changes detected in the ArcC5aR1KO CA1 hippocampal region in one cohort a 7 mo of age (**Figures S6C-F** and (18)).

**Figure 6:**
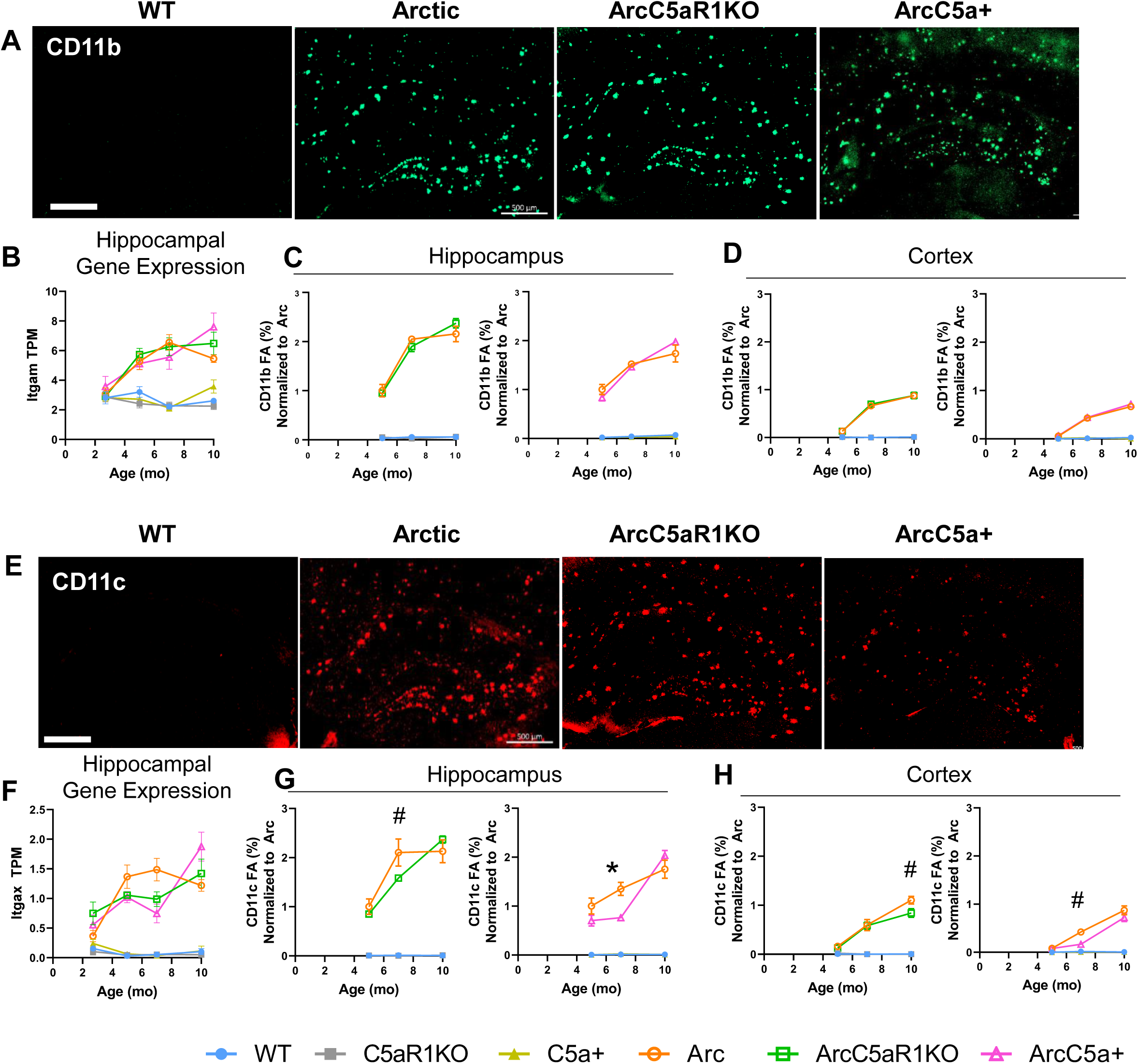
CD11c expression changes with C5a overexpression or C5aR1 ablation. (**A**) representative images from 7 months of CD11b positive microglia in WT, Arctic, ArcticC5aR1KO, and ArcticC5a+ in the hippocampus. (**B-D**) Quantification of gene expression (TPM) of *Itgam* in the hippocampus (**B**) and percent field area of CD11b in hippocampus (**C**) and cortex (**D**) at 5, 7, and 10 months. **E**, representative images from 7 months of CD11c positive microglia in WT, Arctic, ArcticC5aR1KO, and ArcticC5a+ in the hippocampus. (**F-H**) Quantification of gene expression (TPM) of *Itgax* in the hippocampus (**F**) and percent field area of CD11c in hippocampus (**G**) and cortex (**H**) at 5, 7, and 10 months. C5aR1KO and C5a+ cohorts were stained and analyzed separately. Four sections were stained per mouse and the percent field area was averaged over the 4 data points. Data shown as Mean ± SEM. * *p* < 0.05, # 0.1 < *p* < 0.05. Two-way ANOVA with Tukey’s post hoc test. N = 2-6 mice/genotype/age. Scale bar 500 µm.

CD11b is part of the integrin receptor, CR3, with multiple ligands including iC3b/C3b. It is expressed exclusively on microglia in the CNS (56), and previously shown to be upregulated in AD models (57, 58). As expected, CD11b levels were elevated in the hippocampus (**Figure 6A & 6C**), and to a lesser extent in the cortex (**Figure 6D**), of Arc, ArcC5aR1KO, and ArcC5a+ mice compared to their respective controls. CD11b levels peaked at 7 months in the Arc hippocampus, and plateaued thereafter, paralleling our findings from RNAseq data (**Figure 6B**). While in some cohorts CD11b was lower in ArcC5aR1KO at 7 mo, the increase in CD11b was correlated with amyloid load and not downstream complement activation.

CD11c-positive microglia appear early in response to plaque deposition and continue to increase with disease progression around plaques in APP/PS1 mice (59). A time-course of CD11c expression and immunoreactivity (**Figure 6E, 6G-H**) revealed that while essentially absent in the WT, C5aR1KO, and C5a+ genotypes at all ages, CD11c levels increased with age/disease progression in the hippocampus and cortex of Arc, ArcC5aR1KO, and ArcC5a+ mice (**Figures 6G-H**). While CD11c levels were significantly elevated in the Arc hippocampus at 5-10 months compared to controls, we observed 25% and 43% lower CD11c levels in the ArcC5aR1KO and ArcC5a+ hippocampus, respectively, compared to Arc at 7 months (**Figure 6G)**. Furthermore, ArcC5a+ mice had 60% lower CD11c in the cortex at 7 months compared to Arc mice (**Figure 6H**). Interestingly, hippocampal CD11c levels increased to Arc levels by 10 months of age in both ArcC5aR1KO and ArcC5a+. The temporal changes in hippocampal CD11c protein levels in Arc, ArcC5aR1KO, and ArcC5a+ mice match the temporal profile of Itgax gene expression detected with RNA-seq (**Figure 6F**). These findings suggest that blocking C5aR1 may delay or reduce the polarization of a subset of reactive microglia from an early-intermediate stage to a later stage of reactivity in the Arctic model. Thus, appearance of CD11c+ microglia surrounding the plaques may be a key biomarker of microglia that ultimately indicates directly or indirectly cognitive decline and neurodegeneration.

We further assessed pan-reactive and lysosomal markers Iba1 and CD68 (**Figure S7**). Iba1 levels were consistently elevated in Arc, ArcC5aR1KO, and ArcC5a+ hippocampus compared to controls (**Figures S7A & S7D-E**). At 5 months, Iba1 field area was slightly lower in ArcC5a+ compared to Arc, but by 7 months levels were comparable in all three Arc genotypes (**Figure S7D**). CD68 levels were higher in all Arc groups in the hippocampus and to a lesser extent in the cortex compared to WT groups as early as 5 months, matching RNA-seq data (**Figures S7B & SF-H**).

### Astrocyte markers GFAP and C3 are largely unaltered by C5aR1 engagement in the hippocampus

A recent study demonstrated that inflammatory cytokines released by microglia can trigger a shift towards neurotoxic reactivity in astrocytes (19). Thus, to complement the RNA-seq data, we used immunohistochemistry for the pan-reactive astrocyte marker GFAP as well as C3, whose expression has reproducibly been shown to be elevated in neurodegenerative models (19, 60), to characterize changes in astrocyte polarization with disease progression and modulation of C5a-C5aR1 signaling. As expected, we observed a significant increase in GFAP and C3 percent field area in all three Arc groups compared to WT controls in the hippocampus and cortex (**Figure S8A**). Transgene expression of C5a did not significantly change GFAP levels in the CA1 or DG of the hippocampus (**S8C-E**), but did result in decreased levels of GFAP in the CA3 hippocampal region at the early stages of disease (5 mo) relative to Arc mice and a reduction relative to C5aR1-sufficient Arc in cortex at 7mo old age, although the reduction of expression was greater in the C5aR1 deficient Arc cortex at that age (48% reduction) (**Figure S8F**). C3 expression was variable in hippocampal regions with 39% (*p* < .05) decrease in C3 volume in CA1 in ArcC5aR1KO relative to Arc at 7 mo of age (**Figure 7**), while a decrease of 83% of C3 volume and 43% of C3 Field area % were noted in the cortex in two different cohorts of ArcC5aR1KO, relative to Arc mice at 7 months of age (**Figure 7, S8B, S8J**). Lower C3 was seen in the CA1 of ArcC5a+ mice at 7 mo, but, as in ArcC5aR1KO, by 10 mo C3 reactivity was not different from Arc genotype. These data support that ablation of C5aR1, and to a lesser extent C5a overexpression, results in a delay in amyloid-induced astrocyte polarization.

**Figure 7:**
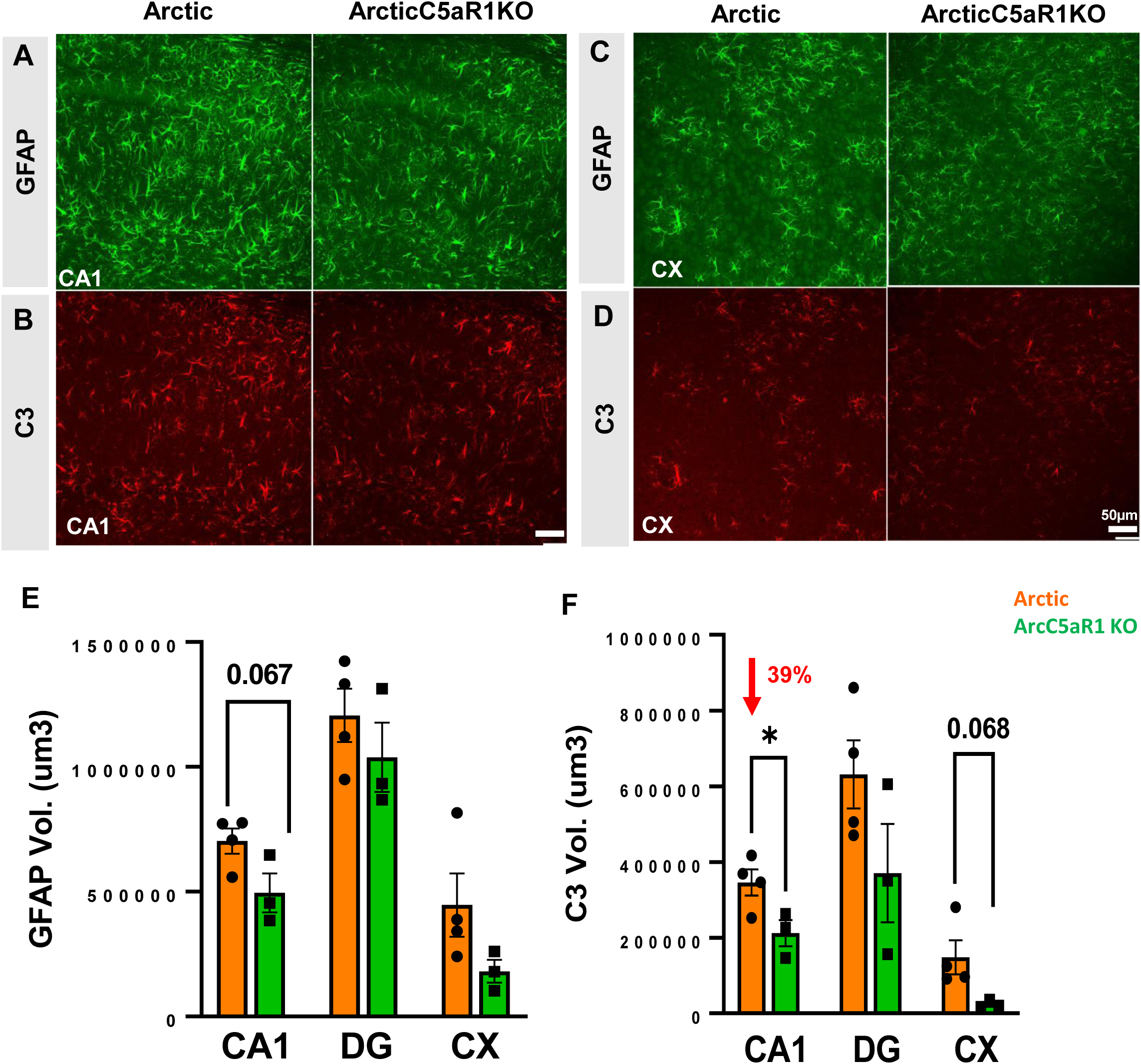
Lower astrocyte C3 expression in ArcticC5aR1KO compared to Arctic mice. Representative confocal images of GFAP immunostaining in CA1 region (A) and in CX (C) region and C3 immunostaining in the CA1 region (B) and in CX region (D) of Arc and ArcC5aR1KO brain at 7 months of age. Quantification of GFAP (E) and C3 (F) volume in the CA1, DG and CX. Z stack (31 steps, 30um) one area/region (confocal Leica SP8, 20x). Data shown as Mean ± SEM. * *p* < 0.05. T-test. N = 3-4 mice/genotype. Scale bar 50 µm.

## Discussion

Using genetic ablation of C5aR1 and transgenic overexpression of the ligand C5a in wild type and Arctic AD model mice, we provide a characterization of cellular, molecular, and transcriptomic changes that occur throughout disease progression and that are modulated by C5aR1 activation in the Arctic mouse model of AD. Further support of a detrimental role of C5a-C5aR1 signaling in the Arctic mouse is shown by the acceleration of hippocampal-dependent spatial memory deficits by overproduction of C5a by a transgene under the GFAP promoter. The upregulated profiles of complement proteins in AD brain (61-63) and the demonstrated safety of the suppression of C5a generation or C5aR1 signaling in human patients with inflammatory diseases (64, 65), suggests C5aR1 antagonism would be well tolerated and may slow cognitive decline in AD and other neurodegenerative diseases in which complement activation is excessive or not appropriately regulated.

For the past decade or so, evidence has been accumulating for a role of the complement system in neuronal injury, synaptic pruning, and cognitive decline (66-69). In the healthy brain, complement proteins tag extranumerary synapses or cellular debris for clearance by microglia (68). However, aberrant complement activity may result in the removal of synapses from stressed neurons in neurodegenerative diseases (70). Complement proteins are upregulated in the human and mouse brain with normal aging and are further increased in AD, models of AD and other neurological diseases (5, 61). Deletion of C1q, C3, or CR3 (CD11b/CD18) reduces microglial engulfment of synapses (68, 71, 72) and protects against memory deficits in aged mice (73) and in models of AD (70, 74) and other neurological disorders (reviewed in (75)). While some have suggested inhibition of upstream complement components (such as C1, C3 and CR3) as a therapeutic strategy, a potential improvement on this strategy would be a more targeted inhibition of the downstream complement activation products, such as the proinflammatory fragment C5a, to preserve the immunoprotective functions of the upstream molecules. For example, C1q can be beneficial by binding to apoptotic neurons to promote clearance by microglia to maintain neuronal homeostasis (76). Previous studies have shown that pharmacologic inhibition or genetic ablation of C5aR1 protects against learning and memory deficits in several mouse models of amyloidosis (9, 18) and that C5a contributes to neuronal injury (7, 8). Landlinger and colleagues provided evidence for a protective effect of an active immunization against C5a in the Tg2576 mouse model of AD, with suppression of CD45^hi^ microglia in immunized mice and some improvement in contextual memory (77). Thus, inhibition of the C5a-C5aR1 pathway could reduce myeloid cell activation as demonstrated here, without interfering with upstream beneficial complement signaling or the formation of the downstream beneficial pathogen killing membrane attack complex (18).

The Arctic48 mutation results in an altered Aβ peptide sequence (E22G) that drives rapid fibril formation and resilience to degradation, resulting in aggressive fibrillar plaque accumulation (78). Using this model, our previous work demonstrated that amyloid is a necessary but not the sole driver of cognitive decline (18), as ablation of C5aR1 prevented loss of neuronal complexity and cognition. Here, our RNA-seq data demonstrated that C5a+- and C5aR1KO-dependent changes in gene expression in the brains of Arctic mice were more prominent in the hippocampus compared to the cortex (with the exception of Rc2) correlating with amyloid load and the significance of hippocampal region on memory and cognition. Eliminating C5aR1 from the Arctic mice did not alter amyloid plaque accumulation, but did delay or reduce expression of genes enriched for inflammatory signaling pathways and microglial cell activation, and several other important AD-associated genes in the hippocampus. Not all AD associated genes were altered, indicating differences in genes whose expression change in response to amyloid plaque deposition and genes whose expression change in response to C5a-C5aR1 signaling, the latter of which may play an essential role in worsening AD cognitive performance. For example, eliminating C5aR1 reduced expression of Inpp5d (SHIP1), which reduces phagocytic capabilities in plaque-associated microglia (45), S100a6, which is found in plaque-associated astrocytes (79), and Stat3, which induces astrocyte reactivity in the 5xFAD model and astrocyte-mediated pro-inflammatory cytokine release in the APP/PS1 model (48, 80), all of which are upregulated and appear to play a detrimental role in human AD brain or mouse models of AD. In addition, the delayed expression of select AD- and DAM1-markers (specifically CD33, Tyrobp, and Trem2) in the C5aR1 deficient Arctic mice suggest a delayed switch from disease mitigating microglia to disease enhancing microglia in mice lacking C5aR1. These transcriptional changes resulting directly or indirectly from C5ar1 ablation coincide with the rescue of cognitive loss (18).

Inflammatory stimuli such as lipopolysaccharide or IFN-γ, or chronic inflammatory states, promote polarization of astrocytes towards a pro-inflammatory state and the subsequent expression and secretion of Lcn2 (81, 82). Furthermore, Lcn2 promotes neuronal loss and hippocampal-dependent memory deficits in a mouse model of vascular dementia (83). While we did not observe C5a-C5aR1 dependent changes in pan-reactive astrocyte GFAP immunoreactivity in the brain, the ArcC5aR1KO group showed delayed expression of a subgroup of pan-reactive and A1 astrocyte genes (Lcn2, Osmr, Cd44, Vim, Serpina3n, Aqp4, C4b and C3), suggesting that while astrocytes do respond to damage associated molecular signals including those resulting from fAβ deposition, there is a delayed activation of neurotoxic astrocytes in the absence of C5aR1. It has been suggested that inflammatory microglia accentuate neurotoxic properties of astrocytes in AD contexts (19), and thus the delayed astrocyte activation seen in the absence of C5aR1 is consistent with delayed secretion of activation signals by microglia (such as TNF, but not C1q). The direct functional consequences of decreased gene expression of neurotoxic astrocyte markers, such as Lcn2, remain to be investigated. However, these findings demonstrate a complexity of astrocyte response mediators that are induced by different insults/injury and are in line with the fine tuning of immune modulating functions that the complement system exerts in multiple other challenges to the host (84-87).

Although overexpression of C5a under the GFAP promoter resulted in acceleration of behavioral deficits in the Arctic mouse, RNA-seq data revealed that overexpression of C5a in the Arctic mice delayed the rise of many inflammatory genes until later in disease progression (10 months). Interestingly, genes associated with synaptic transmission were upregulated in both C5a+ and ArcticC5a+ hippocampi. While these data appear to be opposed to the C5a-induced neuronal damage previously reported, the difference may be due to the in vitro cell cultures and/or the specific in vivo injury (7, 8). Furthermore, the behavioral and transcriptional consequences of C5a overexpression in the ArcC5a+ mice suggest that in addition to C5aR1 signaling, other biochemical and cellular pathways are being induced. Indeed, C5aR2, also referred to as C5L2 or GPR77, is a second known receptor for C5a hypothesized to be a scavenger receptor for C5a and that promotes anti-inflammatory properties (reviewed in (88)). C5aR2 has been shown to be expressed alongside C5aR1 but at much lower levels in bone marrow, myeloid cells, and in the brain. In vitro studies have confirmed that C5a binds to C5aR2 with less affinity than C5a binding to C5aR1, while the rapidly cleaved C5a fragment C5a des Arg binds to C5aR2 with a 10-fold higher affinity than to C5aR1 (89). Formation of β-arrestin-C5aR2 complex results in downregulation of C5a-induced ERK signaling and reduces LPS-induced inflammation (90). Furthermore, C5aR2 can interact with C5aR1, promoting C5aR1 internalization via recruitment of β-arrestin and thus again downregulation of ERK signaling (88).Thus, C5aR2 is also a negative regulator of C5a-C5aR1 activity. C5aR2 agonism reduces C5a-C5aR1-mediated inflammatory response to toll-like receptors, c-type lectin receptors, or cytosolic DNA sensor stimulation of IFN genes (91). Therefore, the reduced expression of inflammatory genes seen in the C5a+ and ArcC5a+ cohorts at 5 and 7 months might be due to the supra-normal levels of C5a derived from the artificial transgene (**Figure S1**) and its subsequent binding to C5aR2. Thus, the anti-inflammatory gene expression in the presence of high concentrations of C5a, emphasizes the therapeutic advantage of the inhibition of C5aR1, without interfering with C5a-C5aR2 engagement as an optimal efficacious strategy for modulating the consequences of C5a generation in AD and perhaps AD related dementias.

C5a overexpression also induced genes involved in lipids and cholesterol metabolic processes, some of which were reduced upon C5ar1 deletion in the Arctic mouse and further enhanced (at later ages) when C5a is transgenically expressed. Although cholesterol cannot cross the blood brain barrier, high levels of circulating cholesterol have been correlated with increased risk of dementia and drugs that reduced levels of cholesterol have been shown to reduce prevalence of AD (92, 93). Our results suggest that cholesterol metabolism may be influenced by C5a-C5aR1 signaling and that C5aR1 inhibition may improve brain cholesterol balance.

Consistent with RNA-seq gene expression, immunohistochemistry showed markedly induced levels of CD11b and CD68 in all Arctic groups at 5-10 mo in hippocampus and cortex. There is evidence to support that microglia high in CD11b in AD may be neuroprotective as well as detrimental. Microglia surround Aβ plaques and promote elimination of deleterious proteins (94), and thus, the early increase in CD11b high microglia in both Arc and ArcC5R1KO mice may be indicative of a microglial disease-mitigating DAM response to amyloid accumulation in the Arc model.

CD11c was also increased in all Arctic mice compared to WT, but at 7 months, the increase was not as pronounced in the ArcC5aR1KO or ArcC5a+ hippocampi compared to Arctic. By 10 months, CD11c levels were comparable between all Arctic groups, suggesting a delay, but not a complete attenuation of differentiation/polarization of this microglial state or subset activation. This early delay in CD11c-positive microglia in the hippocampus of Arc mice lacking C5aR1 may result in the delay of neuroinflammatory pathways that would otherwise contribute to neurodegeneration, providing a possible mechanism of the neuroprotective effects of C5aR1KO. However, continued plaque accumulation as occurs in the Arctic mice due to the fibril inducing mutation may lead to a “tipping point” where C5a-C5aR1 modulation is no longer protective. Transcriptional characterization of CD11c-positive microglia in the APP/PS1 AD model showed that these cells express higher levels of genes associated with cell adhesion, migration, phagocytosis, and lipid/cholesterol processes (59). CD11c was specifically found on microglia that were in contact with amyloid plaques similar to C5aR1 expression, suggesting that these cells are responding to deposited fibrillar amyloid and may play an important role in the immune response to AD pathology (59). Single cell RNA-seq may be able to further functionally define microglia subtypes based on C5aR1, CD11c, and other markers. Of note, since the upregulation of C5aR1 occurs largely in microglia near the plaques, with a paucity of C5aR1 staining of microglia distant from the plaques (20), blocking C5aR1 may have limited or no consequences on homeostatic microglial functions.

## Conclusions

Eliminating C5aR1 from the Arctic mice delayed or reduced expression of genes enriched for inflammatory signaling pathways and microglial and astrocyte cell activation. Given the lack of altered amyloid plaque accumulation in the ArcC5aR1KO mice, these results demonstrate a separation between genes whose expression change in response to amyloid plaque deposition and genes whose expression change in response to C5a-C5aR1 signaling and thus inflammation. Ablation of C5ar1 in Arctic mice also delayed upregulation of complement components, including C4b and C3ar1, suggesting that targeted inhibition of C5aR1 may directly or indirectly (via reduced inflammatory and neurotoxic environment) limit the supply of upstream complement components, such as C4b, which play a role in synaptic pruning as well as microglial activation. Taken together, these data are consistent with a neuroprotective effect of inhibition of the potent anaphylatoxin C5a-C5aR1 interaction, while leaving C1q and C3 intact, making C5a-C5aR1 signaling a promising therapeutic target for AD.

## Abbreviations

AD: Alzheimer’s Disease
ALS: amyotrophic lateral sclerosis
Ast: astrocytes
Aβ: amyloid beta
CP: complement pathway
CPM: count per million
DAM: disease-associated microglia
DI: discrimination index
FDR: false discovery rate
GO: gene ontology
LNA: locked nucleic acid
MAC: membrane attack complex
Mic: microglia
OLM: object location memory
Rc: RNA cluster
TMM: trimmed mean of M-values
TPM: transcripts per million
TSO: template-switching oligo

## Declarations

### Ethics Approval and consent to participate

The Institutional Animal Care and Use Committee of University of California at Irvine approved all animal procedures, and experiments were performed according to the NIH Guide for the Care and Use of laboratory animals.

### Consent for publication

not applicable

### Availability of data and materials

The accession number for the sequencing data reported in this paper is GEO: GSE197591.

### Author Contributions

KC contributed to experimental design, prepared libraries, analyzed RNA-seq data, and was a major contributor in writing the manuscript; NDS contributed to experimental design, prepared libraries, performed IHC experiments, analyzed behavioral and IHC data, and was a major contributor in writing the manuscript; GBG contributed to experimental design, prepared libraries, and assisted with RNA-seq analysis; HYL prepared and sequenced RNA-seq libraries; SHC contributed to experimental design, performed IHC experiments, and analyzed data; PS isolated RNA, performed IHC experiments, analyzed IHC data, AGA performed IHC experiments and analyzed data; TJP performed ELISA experiments and analyzed data; MIF performed IHC experiments and analyzed data; AM contributed to experimental design, data analysis and manuscript preparation; AJT contributed to experimental design, data analysis and manuscript preparation. All authors read and approved the final manuscript

## Acknowledgements

We thank Dr. Scott Barnum (CNine Biosolutions®, LLC) for GFAP-C5a transgenic mice, Dr. Rick Wetsel (McGovern Medical School, University of Texas Health Science Center at Houston) for C5aR1 knock out mice, Dr. Lennart Mucke (Gladstone Institute of Neurological Disease) for Arctic mice, and Dr. Tracy A. Cole for performing and analyzing behavioral testing.

