## Supplemental Figures for "Modulation of C5a-C5aR1 signaling alters the dynamics of AD progression"

### Supplemental Figure 1

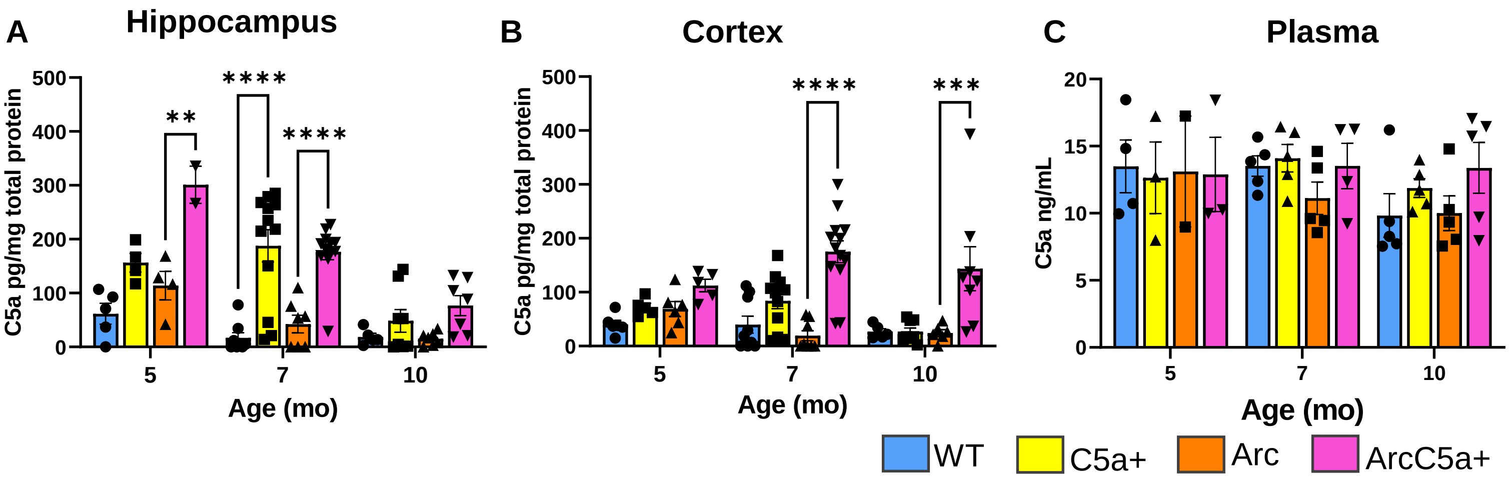


**Figure S1: Presence of the C5aGFAP transgene increases C5a protein levels in brain.** C5a levels detected via enzyme-linked immunosorbent assay (ELISA) as pg/mg total protein hippocampus (**A**) and cortex (**B**), and ng/ml plasma (**C**) in WT, C5a+, Arc, and ArcC5a+ mice at 5, 7, and 10 months of age. Data shown as Mean ± SEM. **$p < 0.01$; *** $p < 0.001$; ****$p<0.0001$. Two-way ANOVA with Tukey's *post hoc* test. N = 2-13 animals per genotype/age for brain. N = 2-5 per genotype/age for plasma.

### Supplemental Figure 2

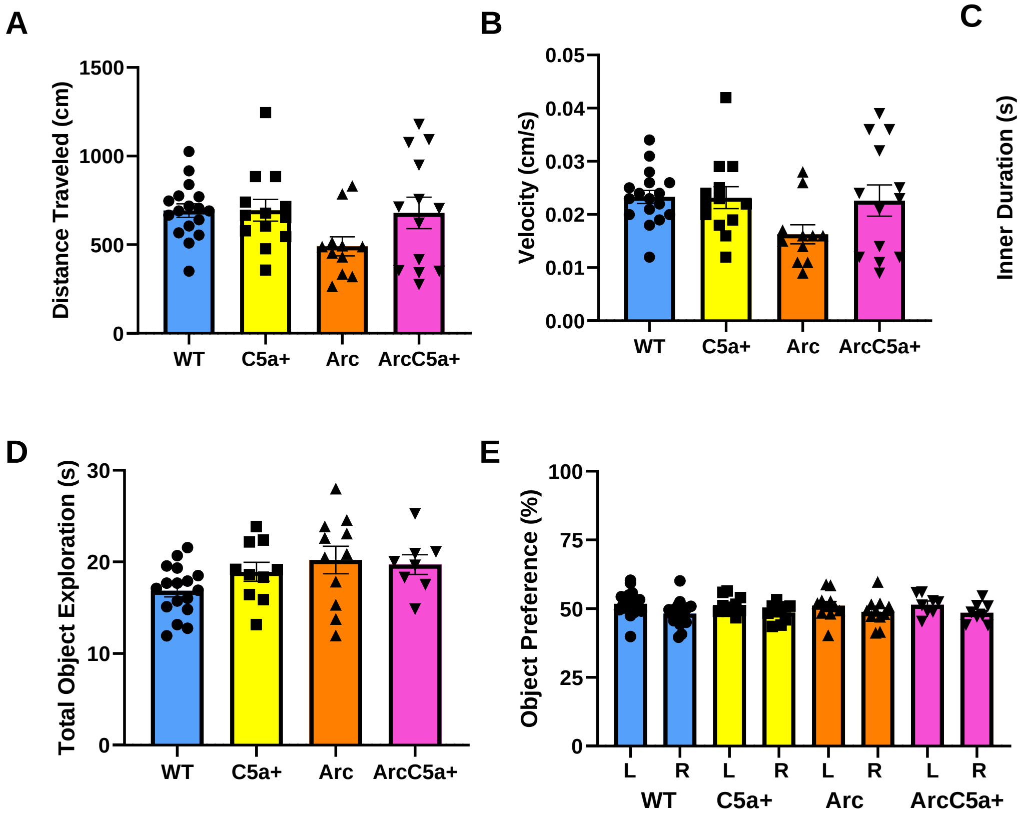


**Figure S2: Locomotion and exploration during open field and object location memory training at 7 months**. (**A-C**) During Open Field (OF), mice were recorded, and distance travelled (**A**), velocity (**B**), and inner duration (**C**) were measured to account for possible changes in locomotion or anxiety-like behaviors. (**D-E**) During object location memory (OLM) training, total object exploration (**D**) and exploration of the left (L) or right (R) objects (**E**) was assessed to eliminate mice that did not sufficiently interact with objects and mice with a bias towards one side. Mice that did not meet training criteria were eliminated from analysis. Data shown as Mean ± SEM. Two-way ANOVA with Tukey's *post hoc* test. Object preference compared with paired t-test. For Open Field N = 17 (WT), 13 (C5a+), 12 (Arc), 13 (ArcC5a+). For OLM training N = 17 (WT), 10 (C5a+), 11 (Arc), 8 (ArcC5a+).

### Supplemental Figure 3

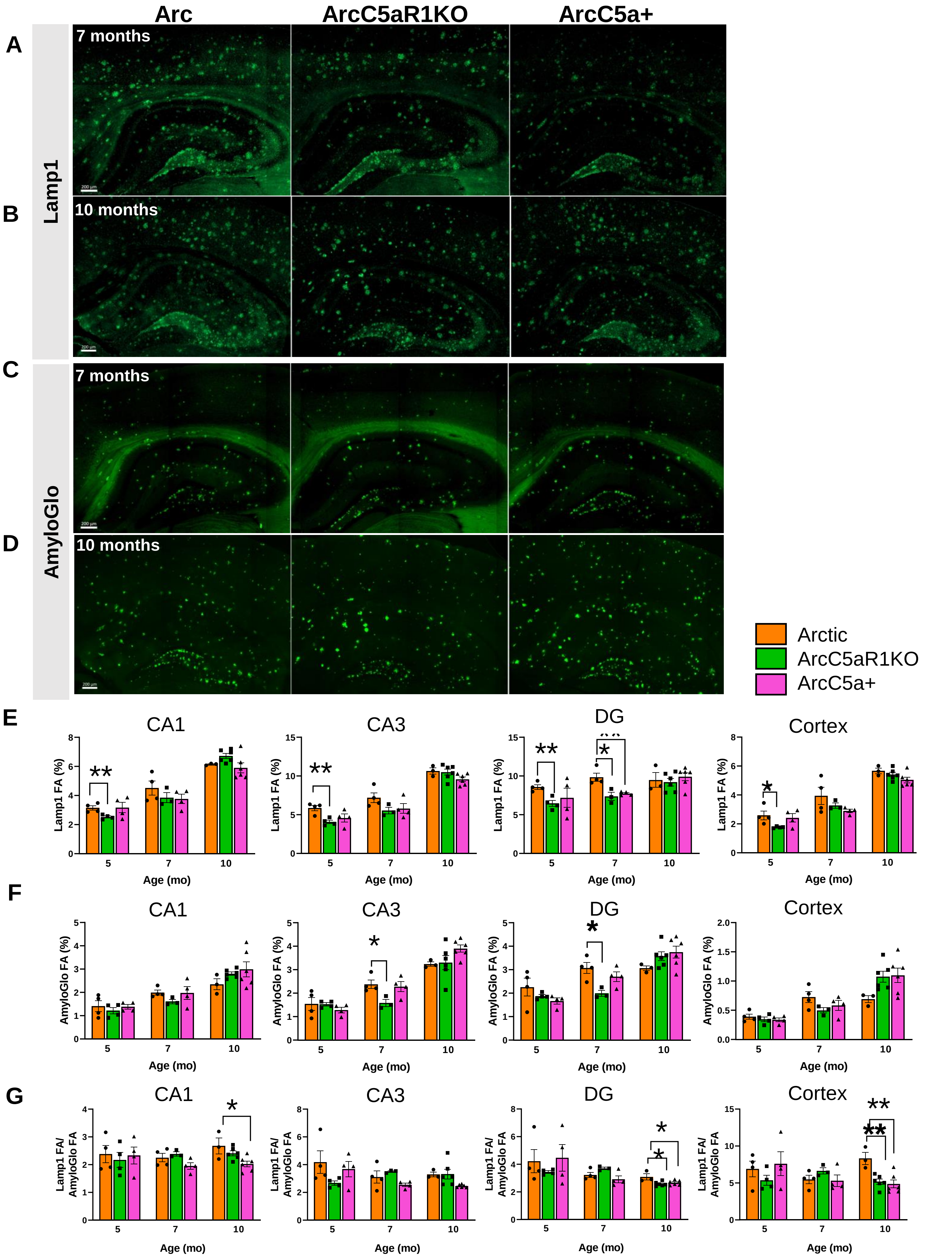

**Figure S3: C5aR1 ablation reduces dystrophic neurites and reduces toxicity of amyloid plaques.** (**A**-**B**) Representative images of Lamp1 immunostaining for Arc, ArcC5aR1KO, and ArcC5a+ mice at 7 and 10 months. (**C-D**) Representative images of AmyloGlo staining for Arc, ArcC5aR1KO and ArcC5a+ mice at 7 and 10 months. Green pseudocolor added to AmyloGlo for visualization (**E**) Quantification of Lamp1 percent field area in CA1, CA3, DG, and cortex. (**F**) Quantification of AmyloGlo percent field area in CA1, CA3, DG, and cortex. (**G**) Comparison of the ration of Lamp1 percent field area to AmyloGlo levels in the CA1, CA3, DG, and cortex. All ages and genotypes were stained and analyzed together. * $p < 0.05$; ** $p < 0.01$. One-way ANOVA with Dunnett's multiple comparisons test. N = 4-6 animals/genotype/age. Three sections were stained per mouse and data was averaged over the three data points. Scale bar 200µm.

### Supplemental Figure 4

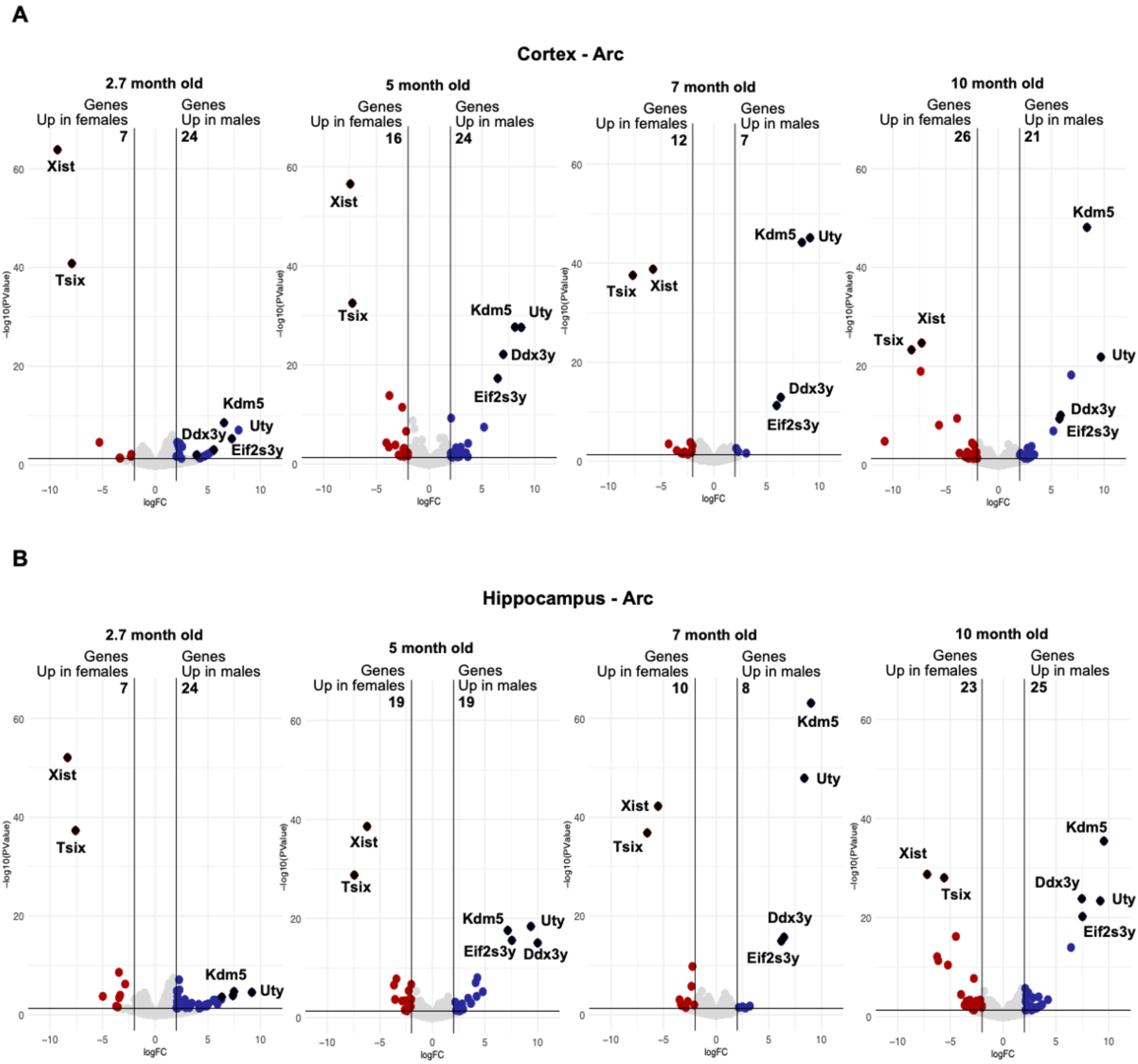


**Figure S4: Differential expression analysis show lack of sex-specific changes in gene expression in the Arctic mice**. (**A**) Volcano plot highlighting genes upregulated in cortices of Arctic female mice (left) and genes upregulated in cortices of Arctic male mice (right) at 2.7, 5, 7 and 10 months of age. Genes upregulated in females and males are colored in red and blue, respectively. p value < 0.05, $|log_2FC| > 2$. Genes highlighted are known sexually dysmorphic genes. (**B**) Volcano plot highlighting genes upregulated in hippocampi of Arctic female mice (left) and genes upregulated in hippocampi of Arctic male mice (right) at 2.7, 5, 7 and 10 months of age. Genes upregulated in females and males are colored in red and blue, respectively. p value < 0.05, $|log_2FC| > 2$. Genes highlighted are known sexually dysmorphic genes. N = 4-10 mice/genotype/age

### Supplemental Figure 5

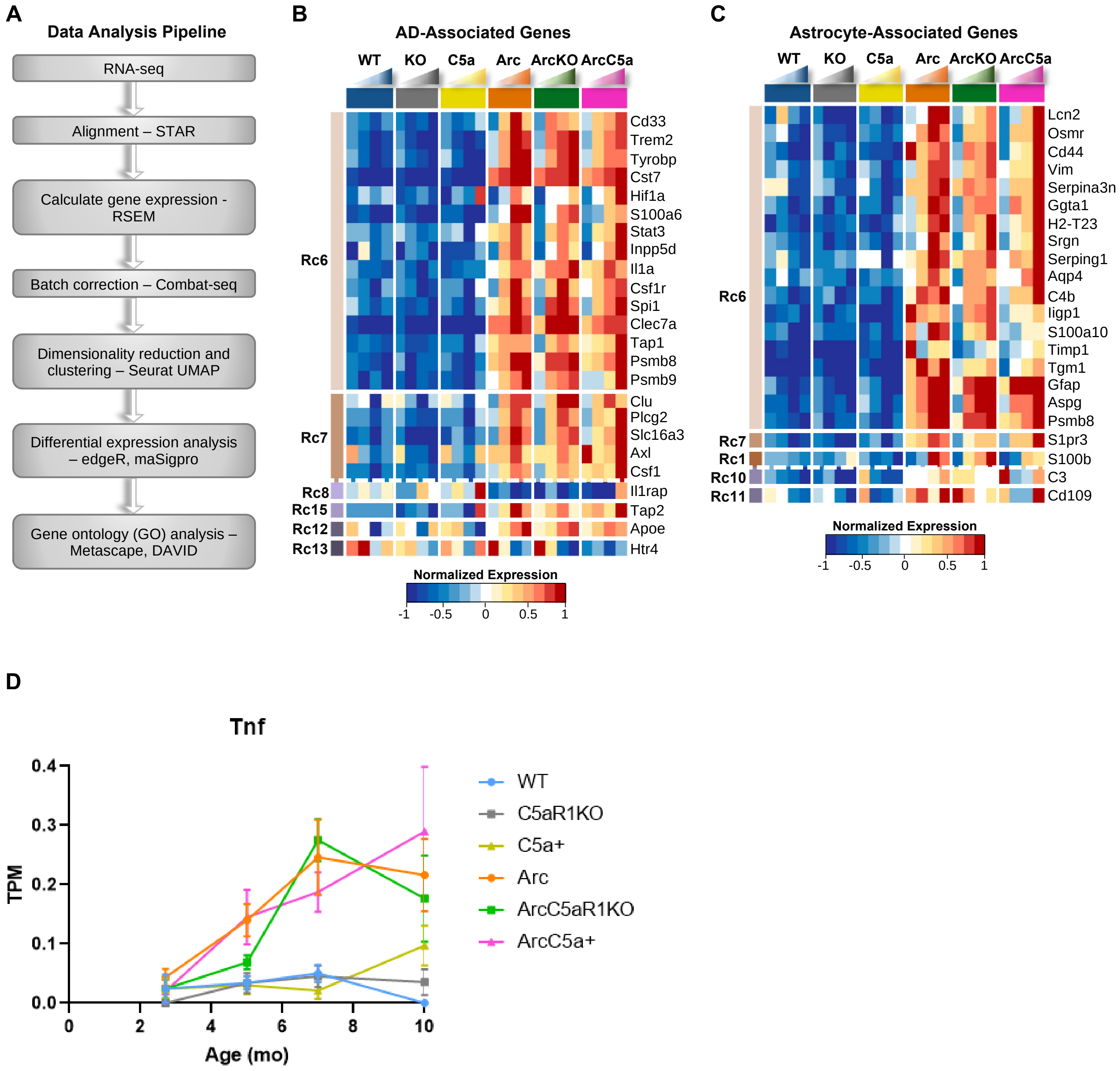


**Figure S5: C5ar1 knockout in Arctic mice delays expression of characteristic AD and astrocyte-associated genes**. (**A**) RNA-seq data analysis pipeline (**B**) Heatmap of 22 genes linked to Alzheimer's disease progression that were present in maSigPro identified clusters. RNA-seq data (TPM) is row-mean normalized. (**C**) Heatmap of 22 astrocyte-associated genes that were present in maSigPro identified clusters. RNA-seq data (TPM) is row-mean normalized. (**D**) Quantification of Tnf expression (TPM) in hippocampus. Data shown as Mean ± SEM. N = 4-10 mice/genotype/age

### Supplemental Figure 6

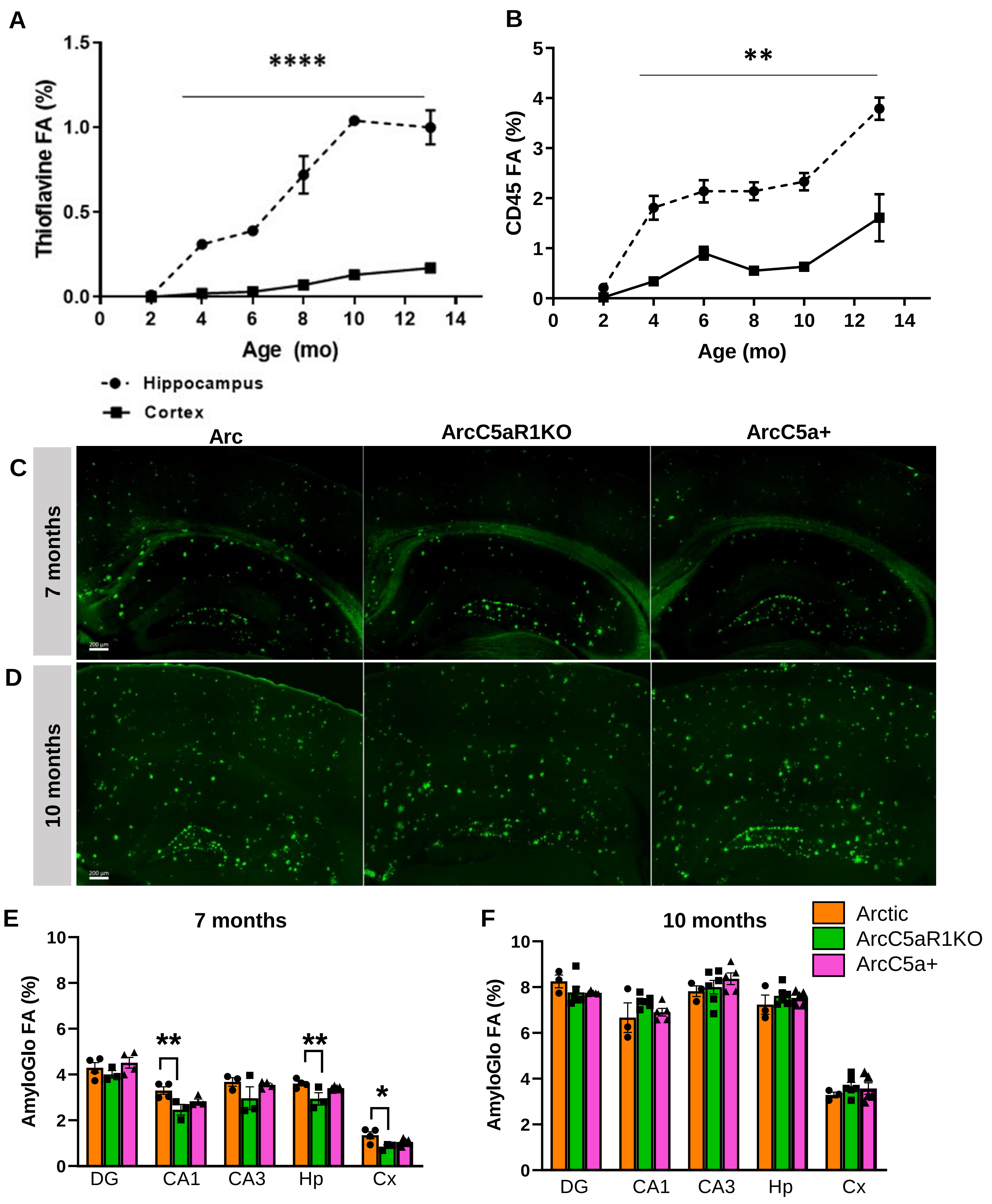

**Figure S6: Plaque accumulation is faster in hippocampus compared to cortex and is unaltered by C5a-C5aR1 modulation.** Time course (2 months to 13 months) of % field area of thioflavin positive plaques (**A**) and CD45 positive microglia (**B**) in the cortex versus hippocampus of Arctic mice. (**C-D**) Representative staining for AmyloGlo in Arc, ArcC5aR1KO, and ArcC5a+ hippocampus at 7 and 10 months along with quantification of percent field area in the hippocampus at 7 months (**E**) and 10 months (**F**). IHC data derived from 4 sections were stained per mouse and the percent field area was averaged over the 4 data points. Data shown as mean ± SEM. *, $p < 0.05$; **, $p < 0.01$; ****, $p < 0.0001$ Two-way ANOVA with Sidak's (**A-B**) or Tukey's (**E-F**) post hoc test. N= 3-6 mice/genotype/age. Scale bar 200 µm.

### Supplemental Figure 7

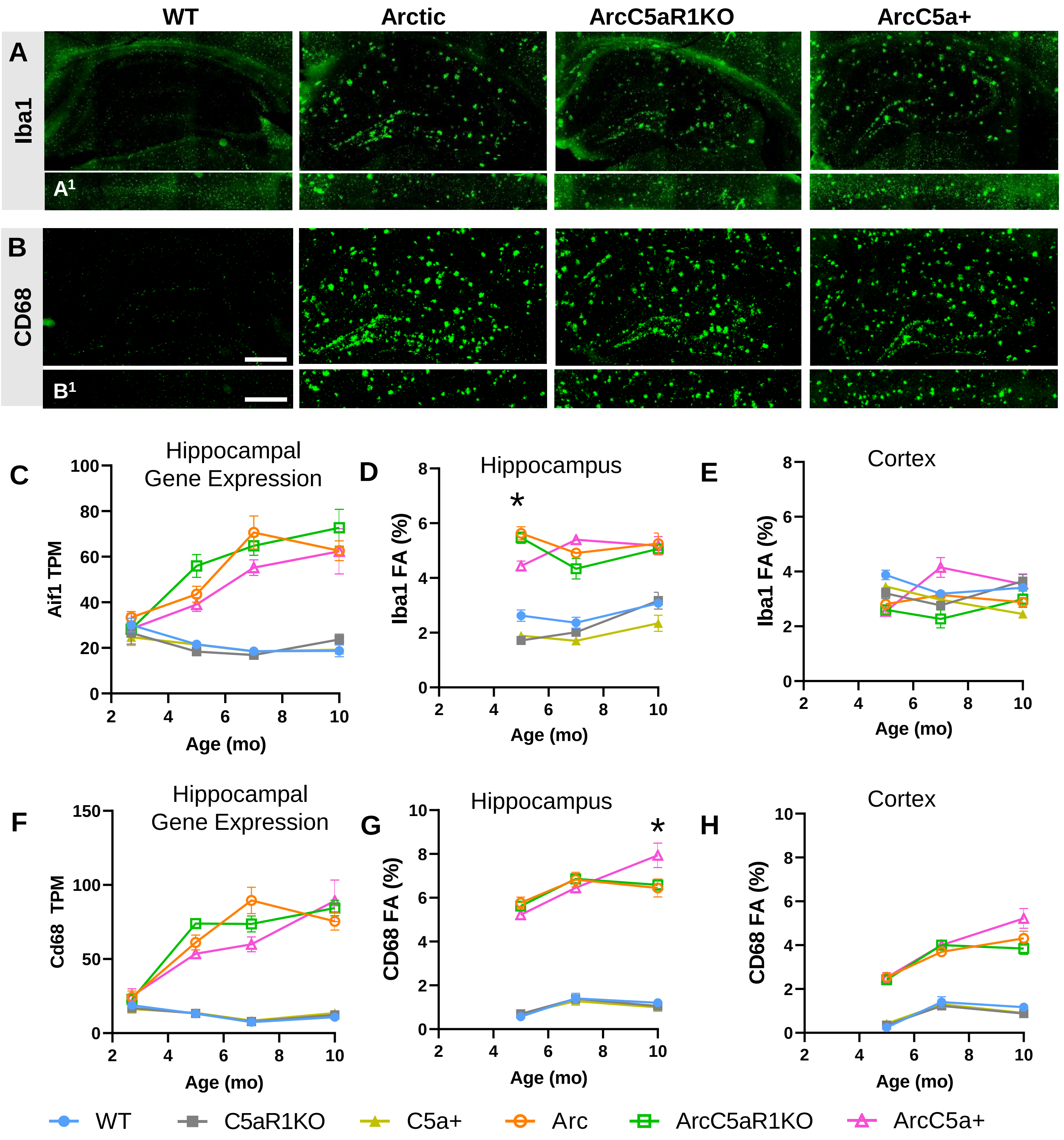

**Figure S7: Iba1 and CD68 not altered by C5a-C5aR1 modulation in Arctic mice, mirroring gene expression.** Representative images from 7 months of Iba1 (**A**) and CD68 (**B**) positive microglia in the hippocampus and cortex (**$A^1$, $B^1$**). Quantification of gene expression (TPM) of Aif1 in the hippocampus (**C**) and percent field area of Iba1 in hippocampus (**D**) and cortex (**E**). Quantification of gene expression (TPM) of CD68 in the hippocampus (**F**) and percent field area of CD68 in hippocampus (**G**) and cortex (**H**). All ages and genotypes were stained and analyzed together**.** IHC data derived from 4 sections stained per mouse and the percent field area was averaged over the 4 data points. Data shown as Mean ± SEM. * $p < 0.05$ Arc vs ArcC5a+. Two-way ANOVA with Tukey's post hoc test. N = 2-6 mice/genotype/age. Scale bar 500 µm.

### Supplemental Figure 8

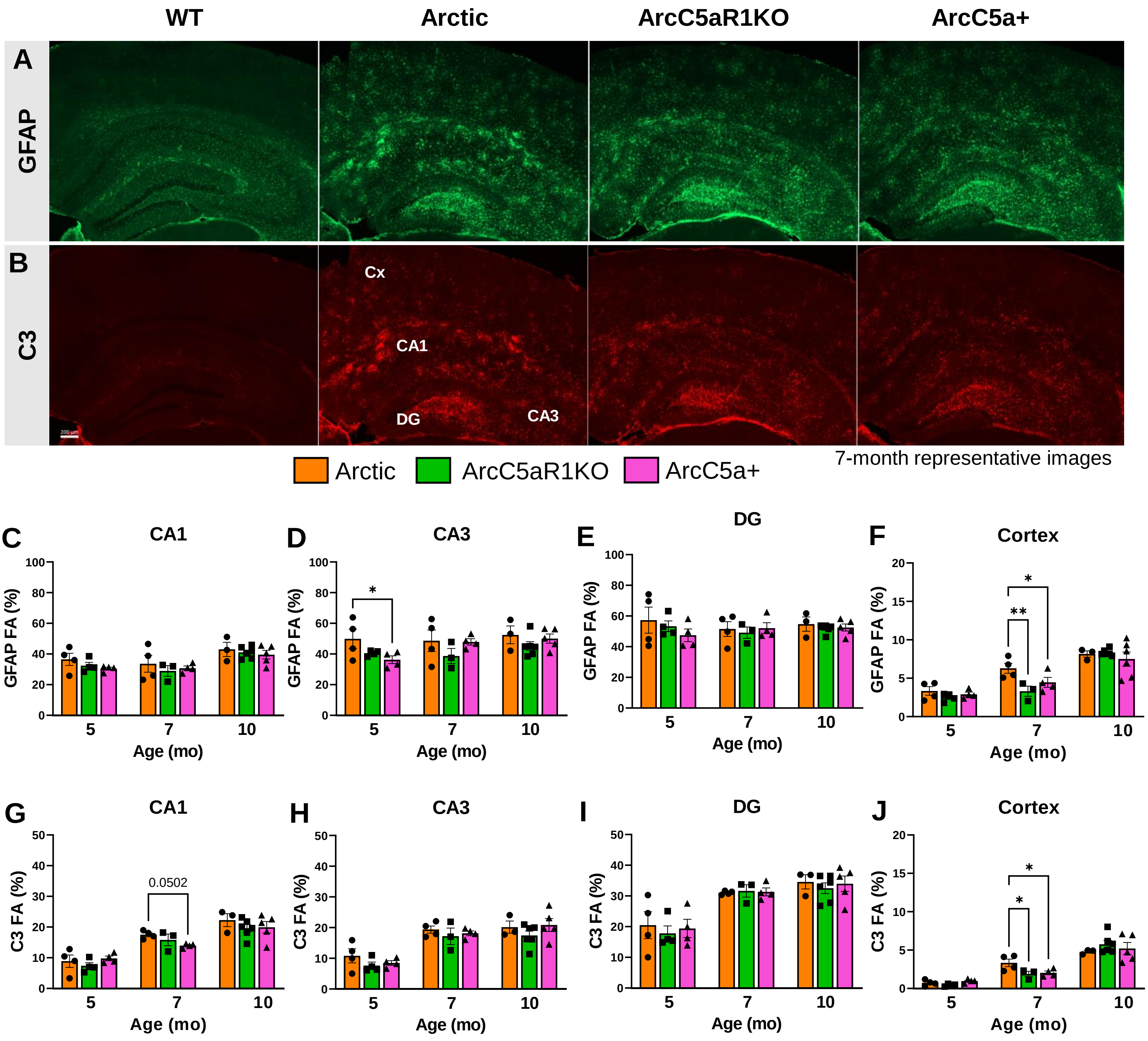


**Figure S8: Astrocyte polarization state in Arctic, ArcticC5aR1KO, and ArcticC5a+ mice.** (A-B) Representative images of immunoreactive staining of GFAP-positive (A) and C3-positive (B) astrocytes in WT, Arc, ArcC5aR1KO, and ArcC5a+ hippocampus and cortex at 7 months of age. Quantification of GFAP (C-F) and C3 (G-J) percent field area (FA) in the hippocampal regions CA1, CA3, dentate gyrus (DG), and in the cortex (Cx) at 5, 7, and 10 months. Four sections were stained per mouse and the percent field area was averaged over the 4 data points. Different ages were stained and analyzed separately. Data shown is Mean ± SEM. * $p < 0.05$. One-way ANOVA with Tukey's post hoc test. N = 3-6 mice/genotype/age. Scale bar 200 µm.
